# Across-breed analyses of genome-wide association studies for stature and mammary gland morphology in cattle reveal pleiotropic effects of the Friesian POLLED haplotype

**DOI:** 10.1101/2025.10.08.681232

**Authors:** Natasha Watson, Qiongyu He, Naveen Kumar Kadri, Alexander S. Leonard, Franz R Seefried, Hubert Pausch

## Abstract

**Background:** Genome-wide association studies (GWAS) in cattle populations have traditionally relied on progeny-derived phenotypes such as estimated breeding values (EBVs) as input phenotypes to identify additive quantitative trait loci (QTL) for complex traits. Increasing availability of cow genotype data now enables GWAS using own performance records to detect both additive and non-additive QTL.

**Results:** Sequence-variant genotypes were imputed for 57,863 cows from the Holstein, Brown Swiss, Original Braunvieh, and Simmental cattle populations that had own performance records for stature and three mammary gland morphology traits (fore udder position, central ligament, front teat position). Genomic heritability ranged from 0.25 to 0.33 for fore udder position, 0.27 to 0.43 for udder central ligament, 0.49 to 0.59 for front teat position and 0.61 to 0.73 for stature. Additive genetic effects explained most of the SNP-based heritability for all traits and breeds. Within-breed genome-wide association studies identified 118 additive and 29 non-additive QTL for the four traits. Non-additive associations were only detected for stature. Although the majority of lead variants were in non-coding regions, we prioritized four missense variants in *HMGA2* (rs385670251), *ZBTB20* (rs470925681), *ARSI* (rs447362502) and *CRAMP1* (rs445465383) as plausible causal variants for four stature QTL. Meta-analysis of the additive GWAS identified 63 mammary gland morphology and 43 stature QTL. A Holstein-specific mammary gland morphology QTL (Chr1:2748715, p=7.59e-35) colocalized with the POLLED locus on chromosome 1. Fine-mapping of this region revealed undesired effects of the Friesian POLLED haplotype on mammary gland morphology traits.

**Conclusion:** Direct phenotypes for a large cohort of genotyped cows provide high statistical power for additive and non-additive association testing. Sequence-based association studies revealed QTL and candidate causal variants for stature and mammary gland morphology traits. Pleiotropic effects of the Friesian POLLED haplotype highlight the need for careful monitoring of potential unintended consequences when selecting for polledness in cattle.

## Background

Genotyping in dairy cattle populations has historically focused on bulls, whereas phenotypic data for economically relevant traits are collected from cows. The bulls’ estimated breeding values (EBVs), de-regressed proofs (DRP), or daughter yield deviations (DYDs) have often been used as pseudo-phenotypes in genome-wide association studies (GWAS) to identify single nucleotide polymorphisms (SNPs) associated with complex traits [1–3]. However, an accumulation of family information in progeny-derived phenotypes can introduce bias in GWAS [4–6] and obscure certain genetic effects, although strategies have been proposed to mitigate some of these biases [7].

As the number of genotyped females increases, the statistical power to analyse genetic associations using their own performance records also grows. Own performance records provide direct phenotypic measurements, allowing for a more accurate quantification of the contributions of both additive and non-additive effects to trait variability [8, 9]. In particular, they enable the estimation of dominance and epistatic effects, which are poorly captured by EBVs or DRPs due to their inherently additive nature [10–12]. Non-additive effects may be particularly relevant at loci associated with inbreeding depression [12].

A range of economically relevant production and conformation characteristics are routinely recorded in dairy cattle. The linear classification of cows during their first lactation provides standardised assessments of these traits. These include, amongst others, height measurements describing stature and mammary gland conformation traits describing morphology of the udder and position and shape of the teats. Udder conformation traits are correlated with udder health, mastitis susceptibility and longevity [13, 14], and therefore are important, easy-to-collect phenotypes to improve functional traits that are difficult to measure.

In this study, we utilised own performance records of 57,863 cows from four breeds to investigate the genetic architecture of four conformation traits. We examined stature as a continuous trait (a well-characterised trait that can be easily compared across studies), and three categorical mammary gland morphology traits. Leveraging imputed sequence variant genotypes, we estimated SNP-based heritability, partitioned it across chromosomes, and decomposed it into additive, dominance, and epistatic components. We conduct within-breed and multi-breed meta-GWAS using additive and non-additive models to identify trait-associated variants. We characterise several QTL and reveal pleiotropic effects of the Friesian POLLED haplotype on mammary gland morphology.

## Methods

### Ethics statement

All analyses were performed using routinely recorded phenotypes and genotypes. We did not perform any experiments on animals, so no ethical approval was required.

### Phenotypes

Phenotypes for stature and mammary gland morphology were obtained from the linear description of cows from four *Bos taurus taurus* breeds (Brown Swiss, Original Braunvieh, Holstein, and Simmental) which is routinely performed in the first lactation. We considered height at the sacral bone and the three mammary gland morphology traits fore udder position, front teat position, and udder central ligament for our investigations. Height is a continuous measurement (in cm) whereas the three mammary gland morphology traits are recorded with a score between 1 and 9. A score of 1 indicates a loose fore udder, large distance between the front teats, and a weak central ligament. A score of 9 indicates a tightly attached fore udder, small distance between the front teats, and a strongly pronounced central ligament. Depending on the trait, records were available for 27,535-27,562 Holstein cows, 23,019-23,143 Brown Swiss cows, 4,138-4,262 Original Braunvieh cows, and 2,896 Simmental cows. The cows’ own performance records were fitted using a linear mixed model implemented in R [15] to account for non-genetic factors. The model included age as fixed effects, and expert and farm/year as random effects. The residuals from this model were used as input for heritability estimation and GWAS (phenotypic outliers, those exceeding 5 standard deviations away from the mean, were removed from the data before analyses were carried out).

### Genotypes

Genotypes for 178,845 animals (60,564 BSW, 12,217 OBV, 89,315 HOL, 14,941 SIM, the remainder are crosses: 1444 OB/BSW, 364 HOL/SIM) were obtained using ten SNP arrays comprising between 20k and 777k SNPs. Only a subset of the genotyped animals had phenotypes. The positions of the SNPs corresponded to the ARS-UCD1.2 (bosTau9) assembly [16] of the bovine genome (only autosomal SNPs were considered in our analyses). We employed PLINK (version 1.9) [17] for quality control separately for each breed and array type. Samples and SNPs with more than 10% missing genotypes and SNPs deviating significantly (p < 1e-07) from Hardy-Weinberg proportions were filtered out.

Imputation was conducted separately for two breed groups: the first group included Brown Swiss, Original Braunvieh and their crosses; the second group included Holstein, Simmental, Swiss Fleckvieh, and their crosses. These two breed groups reflect the structure of the dataset, which includes Swiss Fleckvieh animals with both Holstein and Simmental ancestries, and crosses between Brown Swiss and Original Braunvieh that both share a common ancestral population.

Sequence variant genotypes were imputed into the array-typed genotypes with a two-step imputation procedure using Beagle (version 5.4, [18]). In the first step, genotypes of 690,004 and 669,455 autosomal SNPs typed with the Illumina Bovine HD SNP Bead Chip were imputed using a reference panel comprising 1,183 Brown Swiss and Original Braunvieh cattle, and 1,138 Holstein, Simmental and Swiss Fleckvieh cattle, respectively.

In a second step, genotypes for 26,265,006 sequence variants from 607 in-house sequenced Brown Swiss and Original Braunvieh animals [19] and 37,153,418 sequence variants from 1,464 Holstein and Simmental animals selected from the 1000 Bull Genomes Project run 9 [3] were imputed into the previously imputed high-density genotypes of the respective breeds.

Genotypes with an imputation accuracy greater than 0.4 (Beagle’s DR2 value) were retained for analysis.

### Principal component analysis

For each breed, a genomic relationship matrix (GRM) was built using between 14,094,138 and 16,268,592 autosomal SNPs that had minor allele frequency (MAF) greater than 0.5%, using the “-- grm” flag with GCTA (version 1.94.1) [20]. The principal components (PC) of the GRM were obtained with the “--pca” flag in GCTA.

### Genomic heritability estimation

Genome-wide and chromosome-specific GRMs were constructed for each breed. Additive GRMs were constructed with “--make_grm” of the MPH (v0.52.1) software [21] considering all variants with MAF greater than 0.5%. Dominance GRMs were constructed with the “--dom” flag of MPH. First-order interaction GRMs that account for epistatic interactions were constructed with the “-- make_fore” flag using the dominance and additive GRMs as input for additive x additive interactions, additive x dominance interactions, and dominance x dominance interactions.

We used the “--minque” flag of MPH to estimate the genetic (co)variance components and SNP-based heritability (h^2^_SNP_). For the partitioned h^2^ analysis, we fitted multiple GRMs simultaneously using the “--grm_list” flag. Heritability estimation was performed with variance partitioned across: genome-wide additive GRM, dominance GRM and first-order GRM; per chromosome additive GRMs, dominance GRMs and first-order GRMs; per chromosome additive GRMs. For all heritability estimations, we included the top four PCs of the genome-wide additive GRM as covariates to account for hidden relationships.

### Within-breed genome-wide association study

The association between each trait and imputed sequence variant (MAF > 0.5%) was tested for each breed separately assuming either additive or dominance inheritance. Additive GWAS were conducted using the mixed model-based approach implemented with the “mlma”-flag in the GCTA software [20]. The genome-wide additive GRM and the top four PCs of the genome-wide GRM were included in the mixed linear model to account for relatedness and population stratification [22].

For the non-additive association testing, we estimated the dominance effect (d) using a previously proposed genotype coding scheme [12, 23] as 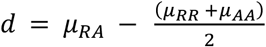, where µ is the phenotypic mean among the three genotype classes for the reference (R) and alternative (A) allele. The genotypes were coded as RR = 0, RA = 2p, and AA = 4p-2, where p is the frequency of the alternate allele. The recoded genotypes were used as input for GCTA (see above) for dominance GWAS.

For additive and dominance GWAS, all variants with a p < 5×10^-9^ were declared significant. This value reflects a Bonferroni-corrected significance threshold for 10 million independent tests, chosen to ensure consistency with the threshold used in the subsequent meta-analyses. The significant variants were then ordered based on their p-values and the most significant variant was retained as top variant, assuming that all other variants within 1 Mb (upstream and downstream) are part of the same QTL and do not constitute independent QTL. Genomic inflation factors were calculated in base R (https://www.r-project.org/).

To perform conditional analysis, we re-ran the mixed linear model and included the expected allele dosage of the top SNP or the POLLED genotype as a covariate. This was carried out within the same GCTA-MLMA framework as the additive GWAS.

### Across-breed meta-analysis of the additive genome-wide association studies

Since the MR-MEGA models require variants to overlap between at least three cohorts (K ≥ 3), 10,119,947 variants were considered for the across-breed meta-analysis using a meta-regression approach [24]. The summary statistics of the additive GWAS were used as input, and the “-qt”-flag was applied for quantitative traits also considering one principal component estimated by MR-MEGA. The “--gco” flag was used for genomic control. SNPs with p-value lower than 5×10^-9^ were declared significant, and QTL were detected as for the within-breed GWAS (see above).

### Variant annotation

Functional consequences of variants were predicted with the Variant Effect Predictor (VEP) software [25] using cache files obtained from the Ensembl (version 104) annotation of the bovine reference genome.

Variants significantly associated with height were extracted from Supplementary Table 2 of Bouwman et al. [2]. The reported coordinates based off the UMD3.1 cattle reference genome were lifted over to the ARS-UCD1.2 coordinates, using chain files provided by the UCSC Genome Browser [26] and the liftover plugin [27] to BCFtools v1.22 [28]. We validated the liftover was successful using the fixref plugin to BCFtools with “--mode stats”. We tested for overlap between the lifted over variants and the significant variants identified in this work using BEDtools v2.31.1 [29] closest, noting the minimum distance between SNPs in the two datasets (with distances of 0 indicating an exact overlap of position).

## Results

### The contribution of genetic and non-genetic factors to trait variability

Additive and non-additive contributions of genome-wide SNPs to the variability of stature and three mammary gland morphology traits were estimated using individual-level genotype and phenotype data of 57,863 cows from four breeds (Figure 1). The genome-wide SNP-based heritability (h^2^_SNP_) varied from 0.67 to 0.77 for stature, from 0.51 to 0.52 for front teat position, from 0.33 to 0.44 for udder central ligament, and from 0.35 to 0.38 for fore udder position, indicating that the four traits considered are moderately to highly heritable. The sample size was between five- and ten-fold larger in Holstein (N= 27,535-27,562) and Brown Swiss (N= 23,019-23,143) than in Simmental (N=2,896) and Original Braunvieh (N=4,138-4,262). Heritability in the larger cohorts was estimated with smaller standard errors (Additional File 1 Table S1-S4) for all genetic variance components considered (additive, dominance, and epistatic). Nonetheless, the non-additive components (dominance, and epistatic) had greater standard errors than the additive component, likely attributable to an increased complexity of the underlying statistical model, which requires larger sample sizes to estimate heritability reliably.

**Figure 1.**
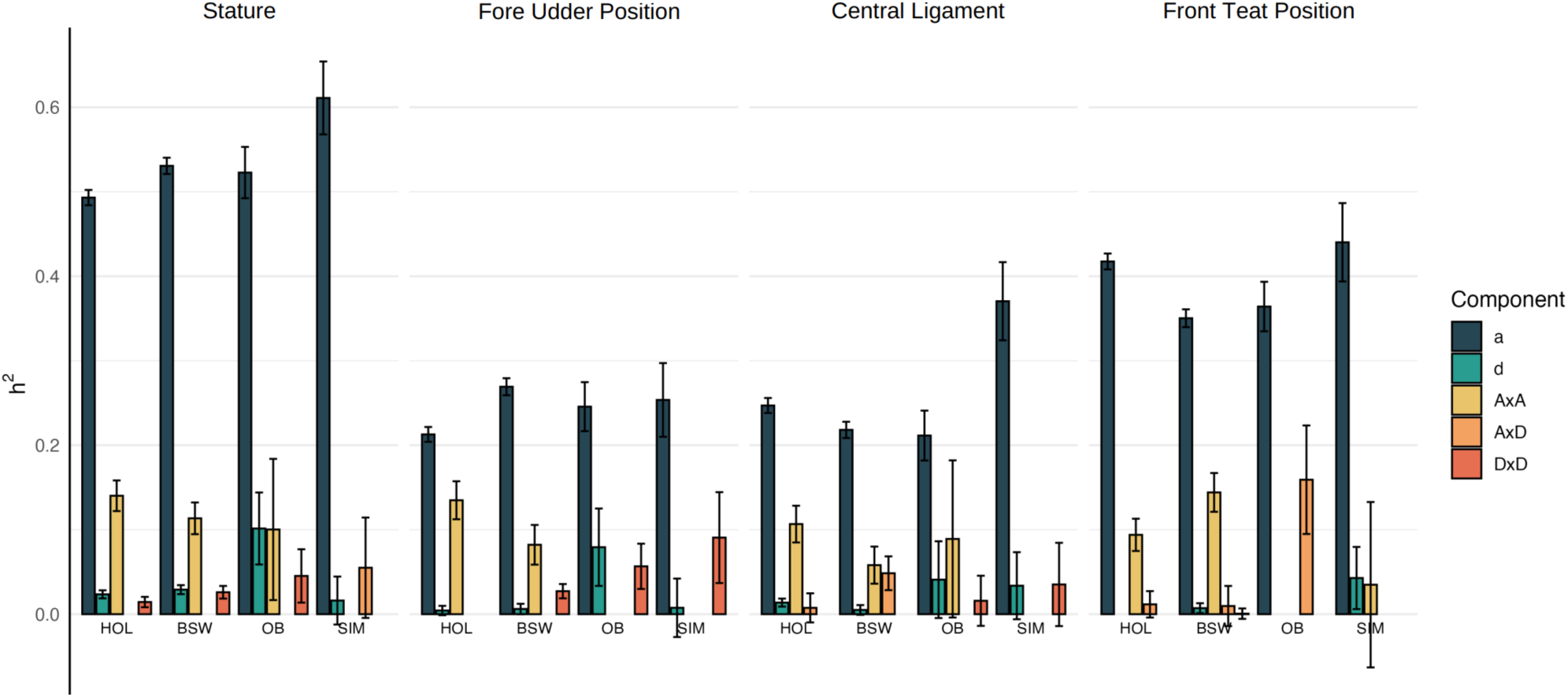
SNP-based heritability (h^2^SNP) for stature (height at sacral bone -KBHM), fore udder position, front teat position and udder central ligament. Breeds included were Brown Swiss (BSW), Original Braunvieh (OB), Holstein (HOL) and Simmental (SIM). A) Genome wide h^2^SNP decomposed into additive (a), dominance (d), additive x additive (AxA), additive x dominance (AxD) and dominance x dominance (DxD) contributions. Error bars represent the standard errors of the estimates. Some confidence intervals contain 0, indicating that the h^2^ estimate is not significantly different from 0.

Additive genetic effects accounted for the largest (between 59-90%) share of the h^2^_SNP_ for all traits and breeds. Additive x additive interactions explained a greater proportion than the other non-additive factors considered (Figure 1). In Holstein and Brown Swiss, for instance, non-additive factors accounted for 20% to 40% of h^2^_SNP_, with additive x additive interactions accounting for 52% to 97% of the non-additive heritability (between 16% to 38% of h^2^_SNP_). Estimates for dominance, additive x dominance and dominance x dominance contributions were small but had relatively large standard errors. Dominance effects were statistically significant only for stature but not for any of the three mammary gland morphology traits.

The relationship between additive h^2^_SNP_ and chromosome length was broadly linear (Additional File 2 Figure S1, Additional File 3 Figure S2, Additional File 1 Table S5). There were notable deviations from this trend (i.e., short chromosomes explaining a large fraction of the heritability (Additional File 1 Table S6)) likely due to chromosomes harbouring loci with disproportionately large effects on the traits considered, such as QTL for stature on BTA25 in Brown Swiss [12] and QTL for mammary gland morphology on BTA17 for several breeds [1, 30]. The observed differences in per-chromosome heritability suggest that the distribution of trait-impacting loci varies between breeds. Some chromosomes contribute disproportionately to the genome-wide h^2^_SNP_ across breeds, while others contribute disproportionately only in some breeds. The former may indicate QTL segregating across breeds while the latter is likely due to breed-specific QTL. Chromosome-level h^2^_SNP_ estimates had greater variability and larger standard errors in the smaller cohorts.

Chromosome-level partitioning of heritability into additive and non-additive components often failed to converge or yielded estimates with substantial standard errors (Additional File 1 Table S4).

### Within-breed single marker GWAS identifies trait-associated variants

We performed genome-wide association studies (GWAS) between imputed sequence variant genotypes and own performance records for stature and mammary gland morphology within each breed using both additive and dominance models. To mitigate confounding effects due to population stratification and relatedness, which can result in an inflation of false positive associations, the statistical models also included the first four principal components of the genomic relationship matrix as fixed effects. The resulting genomic inflation factors ranged from 0.89 to 1.02, indicating that this corrective measure was mostly successful as the test statistics were only slightly in- or deflated.

The additive model identified 18,182 significant variant-trait associations from 16,771 unique variants that defined 60 QTL for stature, 10 for fore udder position, 7 for udder central ligament, and 41 for front teat position across the four breeds studied (Additional File 4 Figure S3, Additional File 5 Figure S4, Additional File 6 Figure S5, Additional File 7 Figure S6 and Additional File 1 Table S7). All significant variant-trait associations from the within-breed GWAS including their functional annotations are provided in Additional File 1 Tables S17-20. We detected more QTL in Holstein and Brown Swiss (50 and 41) than in Original Braunvieh and Simmental (13 and 14).

These discrepancies are most likely attributable to differences in sample size and the resulting statistical power to detect trait-associated loci, rather than to underlying differences in genetic architecture across breeds. We observed several instances, where distinct QTL were located adjacent to one another within and across the breeds. Although most of the significantly associated variants were in intronic or intergenic regions, we also identified 137 highly significantly associated variants overlapping coding sequences that were predicted to have HIGH or MODERATE consequences on the resulting protein. There were only four QTL where variants with HIGH or MODERATE predictions were top or near-top variants. This includes a missense variant of *ZBTB20* with a low (yet tolerated) SIFT score of 0.13 (chr1: 58988339, rs470925681, p.P413L, p=1.32e-15) and a missense variant of *CRAMP1* with a predicted deleterious SIFT score of 0.04 (chr25: 1278638, rs445465383, p.R885Q, p=2.76e-59) that were the lead variants for stature QTL on BTA1 and BTA25, respectively, in Brown Swiss cattle.

The dominance GWAS identified 4,582 significant variants that defined 29 QTL for stature but did not reveal any QTL that reached the genome-wide significant threshold (p < 5×10^-9^) for udder conformation traits (Additional File 8 Figure S7, Additional File 9 Figure S8, Additional File 10 Figure S9, Additional File 11 Figure S10, and Additional File 1 Table S7-S11). These variants were primarily located in non-coding regions of the genome but 3 and 17 were predicted to have HIGH or MODERATE consequences on the resulting protein, respectively.

For Brown Swiss, the significant dominance variants were not significant under the additive model. In the other breeds, the significant dominance variants largely overlapped those detected under the additive model (Additional File 12 Figure S11). For example, a stature QTL located on BTA7 in Holstein (top variant: chr7:61828471, rs482561242, p=8.38e-47) was also identified with the additive model, albeit at much lower significance (Additional File 13 Figure S12). The second top variant at this QTL (chr7:61648112, rs447362502, p=4.83e-41) is a missense variant (p.G303S) in *ARSI* encoding arylsulfatase family member I that has previously been reported as a candidate causal variant underlying a non-additive effect on stature at this QTL [9]. Both top variants are in high linkage disequilibrium (r^2^=0.89) and segregate at moderate minor allele frequency (15%) in the Holstein cattle breed. Neither of the two variants is polymorphic in the other three breeds considered.

The dominance GWAS for stature in Brown Swiss identified three QTL on BTA1, 13 and 25 with top variants residing at 1:58029712 (p=7.30e-10), 13:47208895 (p=5.6e-16) and 25:14475721 (p=1.29e-30), respectively. The top variant at the BTA1 QTL segregated only in Brown Swiss whereas the top variants at the BTA13 and BTA25 QTL segregated at low frequency also in Simmental. None of the top variants segregated in Holstein.

There were instances where the additive and dominance models identified the same genomic region, but the respective lead variants were different. For example, the additive GWAS highlighted a likely deleterious missense variant affecting an evolutionarily highly conserved amino acid residue (chr5:47962581, rs385670251, p.A64P, SIFT score: 0.01, p=1.50e-21, frequency = 0.06, b=-1.8) of *HMGA2* as the top variant for a stature QTL on BTA5 in Simmental. This variant did not segregate in Holstein, Brown Swiss and Original Braunvieh. In contrast, the dominance GWAS in the same population identified a QTL in the same region with an intergenic variant upstream of *HMGA2* as the lead variant (chr5:48000808, rs110936971, p=1.31e-11, frequency = 0.69, b=-0.63). The additive top variant was not significant in the dominance GWAS (p=0.23), while the dominance top variant was nearly ten orders of magnitude less significant (p=7.73e-12) than the top variant from the additive GWAS (Additional File 14 Figure S13). Although the two variants are located within less than 40 kb, they are only in low linkage disequilibrium (r^2^=0.08). Their distinct association patterns and modes of action on stature (Additional File 14 Figure S13) suggest that they represent independent nearby QTL, one (rs385670251) acting additively and the other (rs110936971) non-additively.

### Across-breed meta-analyses detect shared and breed-specific QTL

Meta-analyses of GWAS summary statistics (from the additive model) were performed for the four traits using MR-MEGA. The statistical model implemented in MR-MEGA leverages pairwise allele frequency differences to account for heterogeneity in allelic effects across diverse populations. MR-MEGA was run separately for each trait using 10,119,947 autosomal variants that segregated in at least three populations (Additional File 15 Figure S14). Quantile-quantile plots from the meta-analysis indicate a deviation of the observed test statistic from expectations likely resulting from the large number of QTL identified and extensive linkage disequilibrium (Additional File 16 Figure S15).

The meta-analysis identified 43 and 63 QTL (p < 5×10^-9^) for stature and mammary gland morphology, respectively (Figure 2 and Additional File 1 Tables S12-S15), of which 21 and 36 lead variants showed consistent effects across breeds (Additional File 17 Figure S16). For stature, fore udder position, udder central ligament, and front teat position, we detected 7020, 730, 360, and 4060 significant variant–trait associations, respectively, distributed across 19, 7, 5, and 13 autosomes. All significant variants from the meta-GWAS are provided in Additional File 1 Tables S17-20.

**Figure 2.**
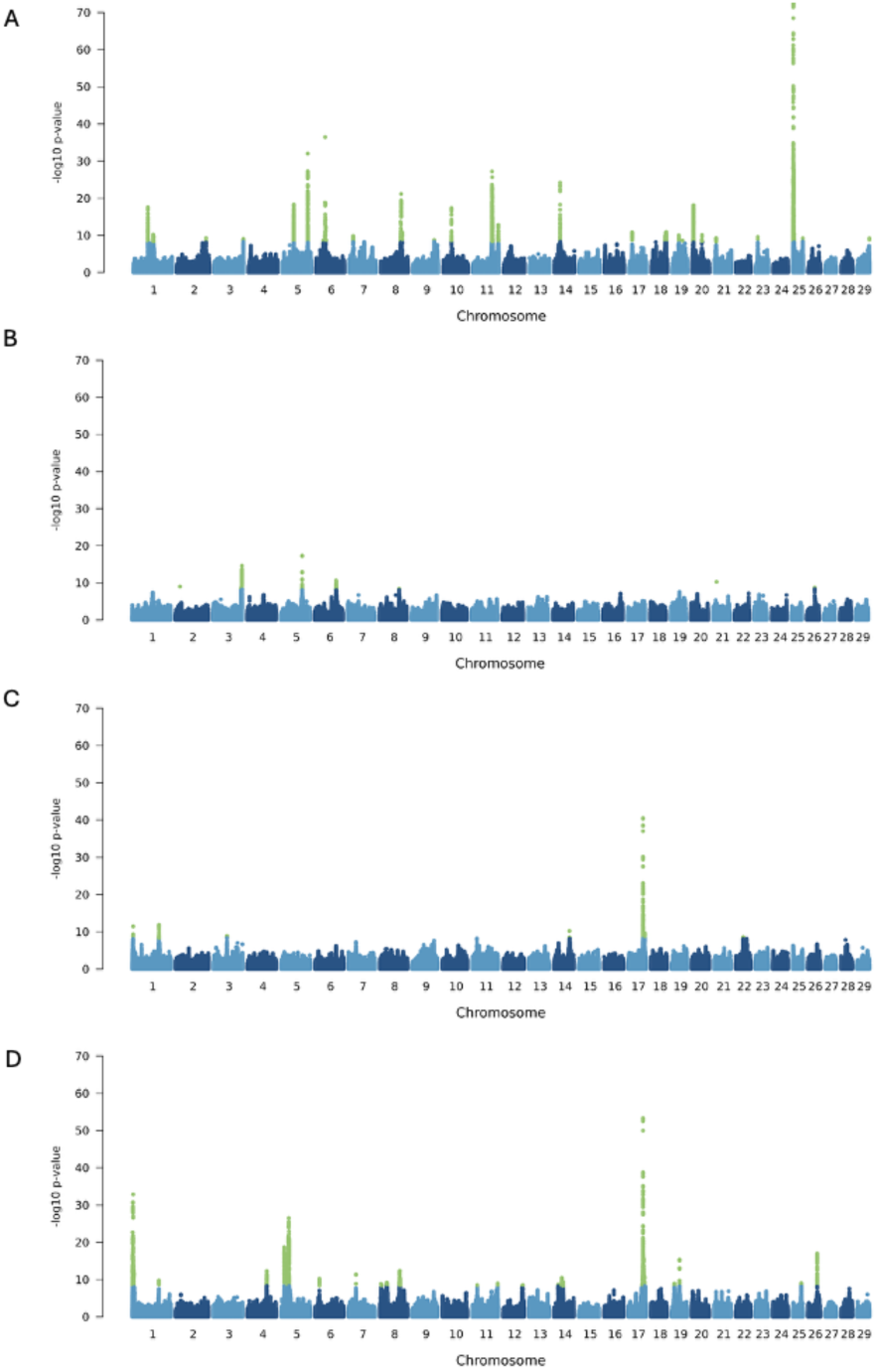
Across-breed meta-analysis of four complex traits. Manhattan plots showing the test statistics for stature (height at sacral bone) (A), fore udder position (B), udder central ligament (C) and front teat position (D). Green points represent SNPs above the significance threshold (p < 5×10^-9^).

We compared the stature QTL identified from the multi-ancestry meta-analysis of the four populations with those that were identified by Bouwman et al. in 17 cattle populations (Additional File 1 Table S16). Of the 43 stature QTL identified, 23 QTL were within 1 Mb from the top SNPs previously identified and an additional 9 QTL were within 2 Mb. Some of these 32 QTL regions overlapped with genes that have been reported to be associated with stature or body size variation in cattle and other species [2], including *PLAG1*, *CCND2*, *LCORL*/*NCAPG*, *HMGA2*, *TCP11*, *STC2*, *INSR2*, *FGFR2*, and *TNNI2*. The remaining QTL identified in our meta-analysis were more than 2 Mb away from any of the QTL reported from Bouwman et al. [2] and might represent distinct associations or capture the effects of multiple nearby QTL. For instance, two QTL on chromosomes 18 and 19 were between multiple QTL identified by Bouwman et al. [2]. Six QTL, which were between 2.2 Mb and 29.1 Mb away from any QTL reported by Bouwman et al. [2] were identified exclusively in the within-breed GWAS of Brown Swiss and/or Original Braunvieh, two breeds not represented in the discovery set of the 1000BG meta-GWAS. This includes the most significantly associated stature QTL from our meta-analysis which resides at the proximal region of BTA25 (Chr25:1485727, p=1.23e-73).

### Characterization of mammary gland morphology QTL

A detailed overview of the most significantly associated variants for the 63 mammary gland morphology QTL is given in Additional File 1 Tables S13-S15. Several signals were shared between central ligament and front teat positioning, including the strongest genome-wide associations that were detected on chromosomes 1 and 17, which likely reflects pleiotropic effects on mammary gland morphology. This is consistent with the well-known influence of the shape of the central ligament on teat positioning. However, no QTL were shared between central ligament and fore udder position or between front teat position and fore udder position. Functional annotation of the lead variants indicated that they all were in non-coding sequence.

The most significant mammary gland morphology QTL on chromosome 17 was associated with both central ligament and front teat position (Figure 3). For central ligament and front teat position there were 371 and 1482 significantly associated markers, respectively, spanning positions 59,658,133 to 67,964,936 bp and 54,882,643 to 70,302,160 bp. An SNP (rs109184112) at Chr17:60,427,314 residing in the intergenic region between *TBX5* encoding T-box transcription factor 5 and *RBM19* encoding RNA binding motif protein 19 was the lead variant for both traits (P_CL_= 2.84e-41, P_FTP_ = 4.53e-54). This QTL (and variant) has previously been implicated with mammary gland morphology in cattle, underscoring the importance of this genomic region across multiple breeds [1, 31]. Homologous regions in the human and mouse genome have also been associated with mammary gland abnormalities [32, 33]. The alternate allele (G) of rs109184112 had a frequency of 98.41% in Holstein, 86.10% in Brown Swiss, 7.68% in Original Braunvieh and 80.17% in Simmental and it showed consistent positive effect directions in the four breeds considered.

**Figure 3:**
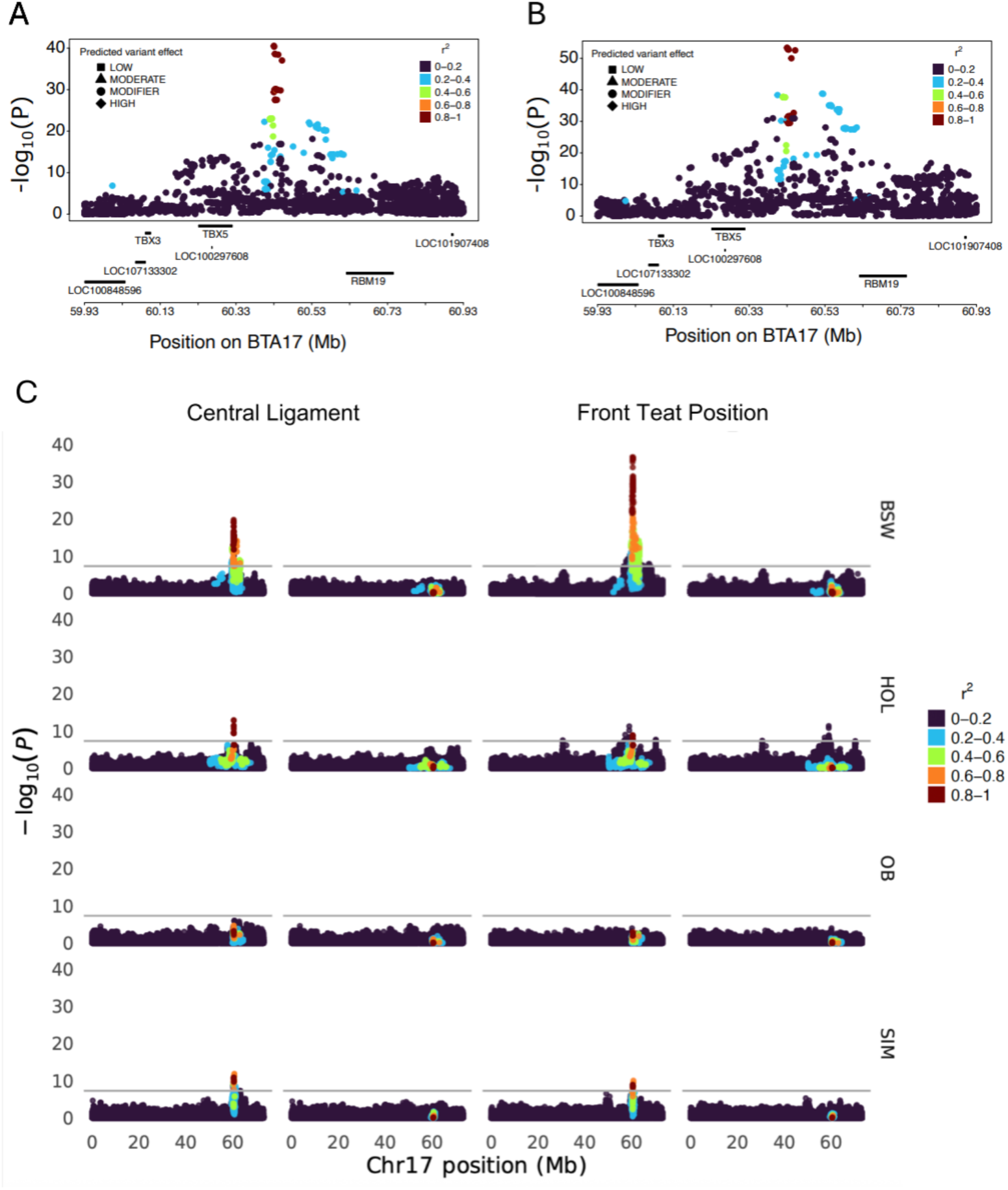
Characterization of a mammary gland morphology QTL on chromosome 17. Zoom plot represents the degree of association of variants at the 60.4 Mb with central ligament (A) and front teat position (B) in the multi-breed meta-analysis. Colour indicates the linkage disequilibrium between in the most significant SNP (Chr17:60,427, 314) from the multi-breed meta-analysis and all other variants. LD was estimated across breeds. C) Manhattan plot representing the degree of association between variants on chromosome 17 with central ligament and front teat position in the within-breed GWAS before (plot on left) and after conditioning (plot on right) the GWAS on the most significant SNP from the meta-analysis (Chr17:60,427, 314). The grey horizontal line represents the significance threshold (p = 5 x 10^-9^).

The mammary gland morphology QTL on BTA17 at 60.4 Mb was absent in the four breeds when the within-breed GWAS for both central ligament and front teat position were conditioned on the top marker identified in the meta-analysis (Figure 3C). However, the conditional association analysis for front teat position in Holstein revealed significant variants (top variant: chr17:58,944,129, p = 4.60e-12) that were more than 1 Mb upstream of the top variant from the meta-analysis, possibly indicating the presence of additional trait-associated variants in this region.

### A mammary gland morphology QTL colocalises with the polled locus

The meta-analysis identified 787 markers on chromosome 1, spanning positions 623,115 to 8,101,744 bp, that were significantly associated with front teat position (p<4.80e-08). The most significant association (p=1.39e-33) was observed at Chr1:3,063,563 (Figure 4a). The minor allele of this variant had a frequency of 2.05% in Brown Swiss, 6.15% in Holstein, 2.43% in Original Braunvieh and 0.59% in Simmental, and it showed consistent negative effect directions in all four breeds. The breed-specific GWAS identified a highly significant QTL (top variant: Chr1:2748715, p=7.59e-35) for mammary gland morphology at the proximal region of BTA1 only in Holstein cattle, with no markers exceeding the significance threshold in other breeds. The top variant (Chr1:3,063,563) from the meta-GWAS was also highly significantly associated (p=3.33e-32) with front teat position in the Holstein within-breed GWAS. Notably, Chr1:3,063,563 also was the top variant for a less significant central ligament QTL (p=3.66e-12).

**Figure 4:**
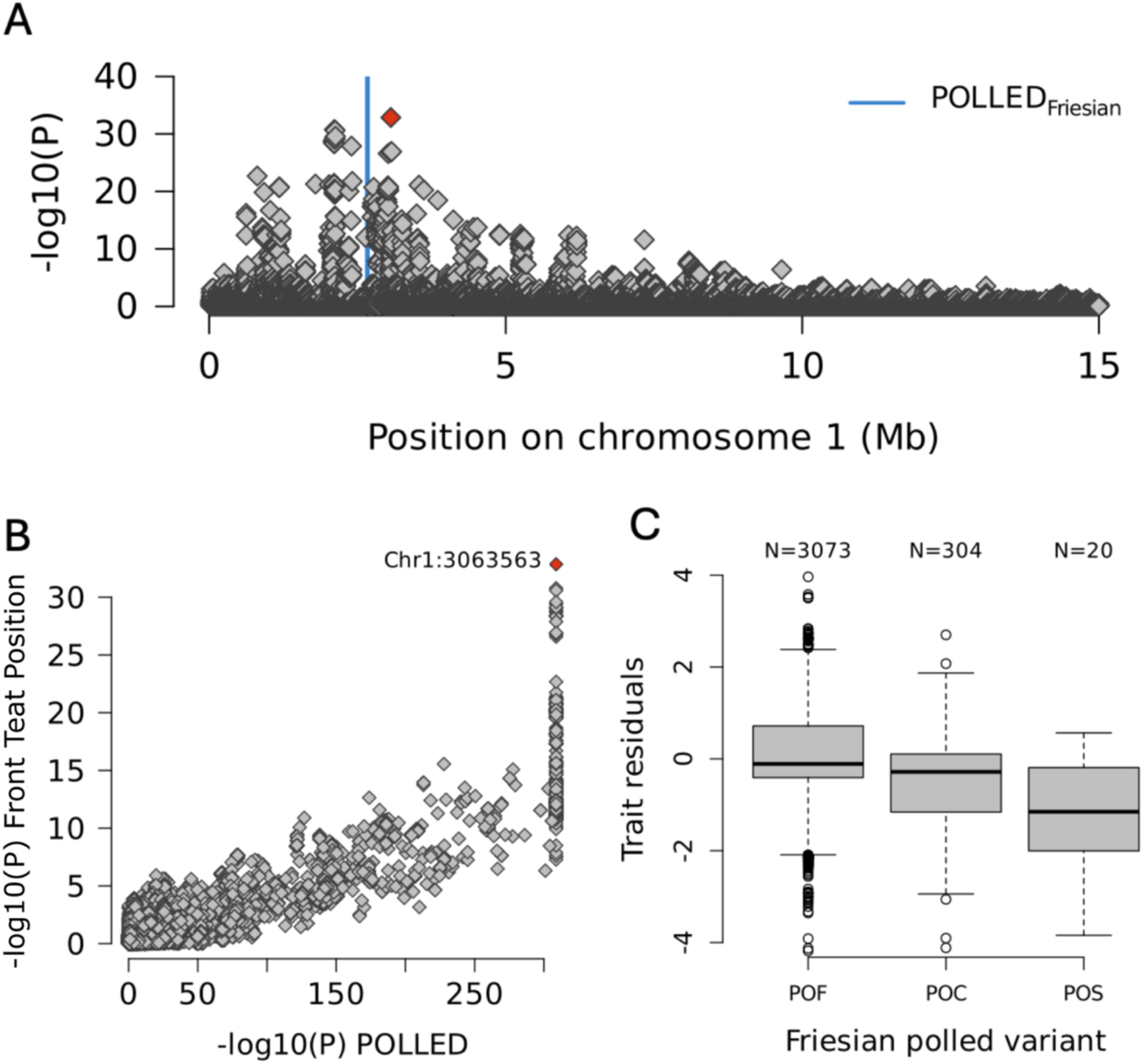
A mammary gland QTL overlaps the POLLED region. A) Manhattan plot representing the degree of association of imputed sequence variants at the beginning of chromosome 1 with front teat position in a multi-breed meta-analysis. Blue colour indicates the position of an 80 kb duplication associated with polledness in Holstein-Friesian cattle. The red symbol represents the most significant SNP (Chr1:3,063,563) from the multi-breed meta-analysis. B) Scatterplot representing the association of imputed sequence variants located at the first 15 Mb on chromosome 1 with POLLED and front teat position. The red symbol represents the most significant SNP (Chr1:3,063,563) from the multi-breed meta-analysis. C) Boxplots representing the association of the Frisian POLLED genotype with front teat position in 3397 Holstein cattle with Axiom array-derived genotypes for the polled allele.

This mammary gland QTL is of particular interest because it overlaps with several alleles associated with polledness in the four breeds studied [34, 35]. In particular, the Friesian polled variant is characterized by an 80 kb duplication on chromosome 1 (g.2629113_2709240dup), which is between the two top variants for the front teat position QTL. The Friesian polled variant has been routinely genotyped in Swiss cattle using the Axiom genotyping array since 2021. Among the 27,562 Holstein cows with front teat position phenotypes, 3,397 had Axiom-derived genotypes available for the Friesian polled variant. Of these, 304 were heterozygous carriers (POC) and 20 were homozygous carriers (POS), corresponding to an allele frequency of 5.06%. We repeated the GWAS for front teat position using this smaller subset of Holstein animals with known polled genotypes. Despite the reduced sample size, the GWAS successfully recovered the QTL on chromosome 1. Although the association signal was attenuated, the Chr1:3,063,563 variant remained highly significant (p=8.39e-11), only slightly less significant than the top marker in this cohort (Chr1:672,192, p=7.97e-12) (Additional File 18 Figure S17a). However, when the GWAS was restricted to 3,073 POF cows—those without the Friesian polled variant—the association with front teat position was completely absent (Additional File 18 Figure S17b). This suggests that the Friesian polled variant or the haplotype containing it, or a variant in high LD with the haplotype, is responsible for the observed association with front teat position.

We repeated the additive GWAS for the four traits in the subset of 3,397 Holstein cattle with known POLLED genotypes with and without fitting the Friesian POLLED genotype as a covariate. As expected, the much smaller sample size resulted in substantially reduced statistical power to detect QTL that were significant at the genome-wide significance threshold. However, the only apparent change we observed between the "normal” and “conditional” analysis across the four traits concerned the mammary gland morphology QTL on BTA1, which was absent when the analysis was conditioned on POLLED. No other QTL were affected by including the POLLED genotype as a covariate, indicating that the POLLED locus does not generally act as a confounder in the dataset.

Association testing between imputed sequence variants and polled genotypes confirmed that the variants most significantly associated with front teat position are also those most strongly associated with polledness (Figure 4b). Furthermore, the Friesian polled genotype exhibited a strong association with front teat position p=3.56e-21, Figure 4c). The effect of the Friesian polled allele on front teat position was substantial, with each copy reducing the trait score by approximately 0.5 points. On average, POF cows (N=3,073) had a front teat position score of 5.26, while POS cows (N=20) scored 3.85, and POC cows (N=304) had intermediate scores (4.71) compatible with an additive effect on the trait. A similar trend was observed for central ligament with POC and POF cows having lower scores than POS cows although the effect of the Friesian polled variant was less pronounced on central ligament than front teat position. These findings indicate that carriers of the Friesian polled variant have more widely spaced front teats and a weaker central ligament.

No mammary gland morphology QTL overlapping the POLLED region was detected in Brown Swiss, although polledness is also strongly selected for in this population. Among the 23,143 Brown Swiss cows with front teat position phenotypes, 1,628 had genotypes available for the Friesian and Celtic polled variants. Of these, 50 were heterozygous carriers (POC) and 4 were homozygous carriers (POS), corresponding to an allele frequency of 1.78%. However, polledness in Brown Swiss is conferred by the Celtic variant which is a 202 bp insertion resulting from the replacement of a 10 bp segment by a duplication of 212 bp. The Friesian polled variant was not detected in the Brown Swiss cohort, consistent with the absence of a front teat position QTL in this breed.

## Discussion

We conducted sequence-based genome-wide association studies for stature and udder conformation in nearly 58k cows from four *Bos taurus taurus* breeds. While several recent association studies have analyzed even larger cohorts of cattle [36–38], our study is among the largest to leverage own performance records, enabling detection of both additive and non-additive genetic effects [39, 40]. The moderate to high heritability of the four investigated traits, together with a relatively large sample size provided high statistical power to identify trait-associated variants with additive effects. In contrast, the power to detect non-additive associations was lower, consistent with the predominant contribution of additive effects to trait variation. As a result, we only detected dominance QTL for stature, which is the trait with the highest overall heritability and the only trait for which the estimated dominance contributions were significant. Larger cohorts are needed to better characterize the non-additive genetic architecture of complex traits [41]. Some dominance QTL overlapped with additive QTL [8, 9, 12], often with stronger significance in the dominance model. For instance, an *ARSI* missense variant (rs447362502, p.G303S) was highly significantly associated for a stature QTL on BTA7 and thus remains a plausible candidate causal variant for affecting height [9]. The dominance GWAS also revealed QTL that were not detected with the additive model. This includes a QTL on BTA25 in BSW cattle which we identified in an earlier study [12]. Markers on chromosomes 7 and 25 had substantial dominance contribution to h^2^_SNP_ in Holstein and Brown Swiss, respectively. Such loci can contribute to inbreeding depression and therefore require further scrutiny [12].

The summary statistics from the additive GWAS were combined across breeds using a multi-population regression framework. Previous multi-breed meta-analyses of stature and mammary gland morphology have been performed [1, 2, 30, 42], but our study is the first to incorporate GWAS results from the Brown Swiss dairy cattle population. This allowed us to confirm previously reported QTL that segregate across multiple breeds including Brown Swiss and detect Brown Swiss-specific QTL. A meta-analysis conducted with imputed sequence variants in 58,265 cattle from 17 populations in the framework of the 1000 bull genomes project identified 163 stature QTL [2]. This study did not include data from Brown Swiss populations in their discovery analysis and missed a large-effect stature QTL that our study detected on the proximal region of BTA25. While even slight methodological differences between studies can result in different outcomes, we consider differences in breed composition to be the most likely explanation for the contrasts observed. Our data suggest that this QTL is specific to the Brown Swiss cattle breed. Although some of the populations included in the 1000 bull genomes meta-analysis are relatively closely related to the Brown Swiss cattle population, this finding shows that even relatively recently diverged populations can carry private variants with large effects on complex traits.

Despite leveraging imputed sequence variant genotypes for a large cohort of cows from four breeds, the identification of causal variants for complex traits remains challenging. Extensive linkage disequilibrium precluded us to readily differentiate between causal variants and variants in linkage disequilibrium with causal variants. Prioritizing candidate causal variants purely based on the statistical significance from GWAS or meta-GWAS can be misleading as the true causal variants aren’t necessarily the most significantly associated variants [43]. Moreover, the meta-analysis approach of MR-MEGA requires that SNPs segregate at least across three of the considered populations. This means that variants specific to one of the considered breeds are filtered out before the meta-analysis. Causal variants for breed-specific QTL are thus represented through linked variants only in the meta-GWAS. By integrating multi- and within-breed GWAS significance with functional annotations, we identified missense variants in *CRAMP1*, *HMGA2*, *ZBTB20*, and *ARSI* as lead variants for stature QTL. Our findings corroborate the previously suggested recessive effect of the *ARSI* p.G303S variant (rs447362502, chr7:61648112) on stature [9]. This variant occurred only in the Holstein population. A relatively rare p.A64P variant of *HMGA2* (rs385670251, chr5:47962581) that only occurred in Simmental was the top variant for an additive stature QTL in Simmental. This variant was also the lead variant for a stature QTL in cattle in a previous meta-analysis conducted by the 1000 Bull Genomes Project consortium [2]. *HMGA2* missense variants are also associated with stature in species other that cattle [44–46]. We also identified an independent dominance QTL for stature upstream *HMGA2* in Simmental cattle further corroborating that this region is important for growth traits in cattle. While the two missense variants in *ZBTB20* and *CRAMP1* have not been reported to be associated with cattle stature yet, the *ZBTB20* gene has previously been reported to harbour alleles with large effects on stature in other species [47, 48]. Collectively, these findings indicate that the identified missense variants represent potential candidate causal variants underlying the observed stature QTL.

A QTL for mammary gland morphology at the proximal region of BTA1 colocalizes with the POLLED locus. This QTL was identified in Holstein cattle, but not in the other breeds investigated. To our knowledge, potential pleiotropic effects of the Friesian polled haplotype on economically relevant traits have not been reported before. Both the Celtic and Friesian POLLED variants are associated with minor atypical eyelash and eyelid phenotypes, indicating that this genomic region may have broader implications on epidermal tissue structures [49]. Although the Celtic and Friesian variants are located near each other and produce the same hornless phenotype, they differ substantially in size. The Friesian variant consists of an 80 kb duplication, while the Celtic variant is a 212 bp indel [34, 35]. This difference in size suggests the potential for more extensive regulatory or functional consequences of the Friesian variant for neighbouring genes, despite the two variants being indistinguishable in terms of polledness. The mammary gland morphology QTL was absent when the association study was conducted in animals not carrying the Friesian POLLED haplotype. This provides strong evidence that the mammary gland morphology QTL is either due to the Friesian POLLED variant itself or due to a variant in high linkage disequilibrium with the Friesian POLLED variant. Fine-mapping efforts in cohorts much larger than ours are required to differentiate between tightly linked candidate variants [50]. Regardless, the observed additive association with a weaker central ligament and increased distance between the front teats in cows carrying the Friesian polled allele support an undesired pleiotropic effect of the Friesian POLLED haplotype. A weak central ligament correlates with reduced productive life in dairy cattle [51].

Given an increasing selection pressure for polledness in breeding programs, our findings underscore the need for careful monitoring of potential unintended consequences associated with POLLED alleles, particularly those affecting mammary gland morphology and other economically important traits. However, because traits such as mammary gland morphology are highly polygenic, selection strategies can likely offset the undesirable effects associated with the Friesian POLLED haplotype.

## Conclusions

Imputed sequence variant genotypes, combined with individual performance records for a large cohort of cows, enabled a detailed investigation of the additive and non-additive genetic architecture of four complex traits. Additive effects explained most of the SNP-based heritability for stature and mammary gland morphology in the four studied cattle populations. Association analyses identified numerous additive QTL, while non-additive QTL were comparatively rare.

Pleiotropic effects of the Friesian POLLED haplotype on mammary gland morphology highlighted the need for careful monitoring of potential unintended consequences associated with strong selection for polledness.

## Declarations

### Ethics approval and consent to participate

Not applicable.

### Consent for publication

Not applicable.

### Availability of data and materials

Genotype and phenotype data analyzed in this study are owned by Swiss cattle breed associations and not publicly available. Summary data from the GWAS are available from the corresponding author upon reasonable request.

## Authors’ contributions

Preparation of genotype and phenotype data: NW NKK QH FRS; GWAS, heritability estimation: NW; QTL fine mapping and interpretation of results: NW ASL HP; Manuscript writing and revision: NW HP. All authors read and approved the final manuscript.

## List of abbreviations

EBV: estimated breed value
GWAS: Genome-wide association study
SNP: Single nucleotide polymorphism
QTL: Quantitative Trait Locus
BSW: Brown Swiss
OB: Original Braunvieh
HOL: Holstein
SIM: Simmental
LD: linkage disequilibrium
MAF: minor allele frequency
GRM: genomic relationship matrix

## Acknowledgements

We thank the Swiss cattle breeding associations for providing pedigree, phenotype and genotype data of cattle.

## Additional files

**Additional file 1 Table S1-S20**

Format: xlsx

Title: Supporting Tables

Description: Excel spreadsheet containing separate tabs for the 20 Supporting Tables. The first tab contains a detailed description of the information given.

**Additional file 2 Figure S1**

Format: tiff

Title: Additive SNP-based heritability (h^2^) per chromosome across four breeds for stature and mammary gland morphology traits.

Description: Additive h^2^ partitioned across chromosomes plotted against chromosome length (Mb). Breeds included were Brown Swiss (BSW), Original Braunvieh (OB), Holstein (HOL) and Simmental (SIM). Traits are stature with height at sacral bone, and mammary gland morphology traits are fore udder position, front teat position and udder central ligament. Vertical lines outside the circles indicate the standard errors. The black line is a regression line, and the grey shaded area represents the 95% confidence interval.

**Additional file 3 Figure S2**

Format: tiff

Title: Dominance SNP-based heritability (h^2^_SNP_) per chromosome for two breeds for stature and mammary gland morphology traits.

Description: Dominance h^2^ (from additive and dominance h^2^ estimation) partitioned across chromosomes, plotted against chromosome length (Mb). Breeds included were Brown Swiss (BSW) and Holstein (HOL). Traits are stature with height at sacral bone (KBHM), and the mammary gland morphology traits are fore udder position, front teat position and udder central ligament. Vertical lines outside the circles indicate the standard errors. The black line is a regression line, and the grey shaded area represents the 95% confidence interval.

**Additional file 4 Figure S3**

Format: pdf

Title: Additive genome-wide association studies for stature across four breeds.

Description: Breeds included were Brown Swiss (BSW), Original Braunvieh (OB), Holstein (HOL) and Simmental (SIM). The –log10(p) values are plotted against the genomic position by chromosome. The horizontal line denotes the genome-wide significance threshold (p < 5×10^-9^).

**Additional file 5 Figure S4**

Format: pdf

Title: Additive genome-wide association studies for fore udder position across four breeds.

Description: Breeds included were Brown Swiss (BSW), Original Braunvieh (OB), Holstein (HOL) and Simmental (SIM). The –log10(p) values are plotted against genomic position by chromosome. The horizontal line denotes the genome-wide significance threshold (p < 5×10^-9^).

**Additional file 6 Figure S5**

Format: pdf

Title: Additive genome-wide association studies for udder central ligament across four breeds.

Description: Breeds included were Brown Swiss (BSW), Original Braunvieh (OB), Holstein (HOL) and Simmental (SIM). The –log10(p) values are plotted against genomic position by chromosome. The horizontal line denotes the genome-wide significance threshold (p < 5×10^-9^).

**Additional file 7 Figure S6**

Format: pdf

Title: Additive genome-wide association studies for front teat position across four breeds. Description: Additive genome-wide association test for udder central ligament across four breeds.

Description: Breeds included were Brown Swiss (BSW), Original Braunvieh (OB), Holstein (HOL) and Simmental (SIM). The –log10(p) values are plotted against genomic position by chromosome. The horizontal line denotes the genome-wide significance threshold (p < 5×10^-9^)

**Additional file 8 Figure S7**

Format: pdf

Title: Dominance genome-wide association studies for stature across four breeds.

Description: Breeds included were Brown Swiss (BSW), Original Braunvieh (OB), Holstein (HOL) and Simmental (SIM). The –log10(p) values are plotted against genomic position by chromosome. The horizontal line denotes the genome-wide significance threshold (p < 5×10^-9^).

**Additional file 9 Figure S8**

Format: pdf

Title: Dominance genome-wide association studies for fore udder position across four breeds.

Description: Breeds included were Brown Swiss (BSW), Original Braunvieh (OB), Holstein (HOL) and Simmental (SIM). The –log10(p) values are plotted against genomic position by chromosome. The horizontal line denotes the genome-wide significance threshold (p < 5×10^-9^).

**Additional file 10 Figure S9**

Format: pdf

Title: Dominance genome-wide association studies for udder central ligament across four breeds.

Description: Breeds included were Brown Swiss (BSW), Original Braunvieh (OB), Holstein (HOL) and Simmental (SIM). The –log10(p) values are plotted against genomic position by chromosome. The horizontal line denotes the genome-wide significance threshold (p < 5×10^-9^).

**Additional file 11 Figure S10**

Format: pdf

Title: Dominance genome-wide association studies for front teat position across four breeds.

Description: Breeds included were Brown Swiss (BSW), Original Braunvieh (OB), Holstein (HOL) and Simmental (SIM). The –log10(p) values are plotted against genomic position by chromosome. The horizontal line denotes the genome-wide significance threshold (p < 5×10^-9^).

**Additional file 12 Figure S11**

Format: pdf

Title: Significant chromosomes dominance vs additive SNPs for stature.

Description: Additive and dominance GWAS results for stature were compared by plotting – log10(p) values from both models for chromosomes showing genome-wide significance under the dominance model. Breeds included were Brown Swiss (BSW), Original Braunvieh (OB), Holstein (HOL) and Simmental (SIM). Variants were classified as significant (p < 5×10^-9^) under the additive model, the dominance model, or both models, and coloured accordingly (additive = dark blue, dominance = teal, both=orange).

**Additional file 13 Figure S12**

Format: pdf

Title: A stature QTL on BTA7 in Holstein cattle.

Description: Effect of chr7:61828471 on stature in Holstein cattle. This marker reached genome-wide significance under the dominance model (p= 8.38e-47). Phenotype distributions are presented for each genotype class.

**Additional file 14 Figure S13**

Format: pdf

Title: A stature QTL on BTA5 in Simmental cattle.

Description: A) Association testing with the additive (grey) and dominant (blue) model indicate the presence of QTL at approximately 48 Mb on bovine chromosome 5 encompassing the *HMGA2* gene. B) Scatterplot comparing the association (-log10(P)) of variants from the additive (ADD) and dominance (DOM) model. C) Boxplots representing the effects of the top variants from the dominance (green) and additive (grey) models on stature.

**Additional file 15 Figure S14**

Format: pdf

Title: Overlap of significant SNPs identified by MR-MEGA meta-analysis and single-breed GWAS.

Description: Number of shared and unique SNPs surpassing the genome-wide significance threshold (p < 5×10^-9^). Breeds included were Brown Swiss (BSW), Original Braunvieh (OB), Holstein (HOL) and Simmental (SIM). Traits included were (A) stature (height at sacral bone - KBHM, (B) fore udder position, (C) udder central ligament and (D) front teat position.

**Additional file 16 Figure S15**

Format: pdf

Title: QQ plots from MR-MEGA meta-analysis.

Description: Traits included were (A) stature (height at sacral bone), (B) fore udder position, (C) udder central ligament and (D) front teat position. The observed -log₁₀(p) values are plotted against the expected distribution under the null hypothesis.

**Additional file 17 Figure S16**

Format: pdf

Title: Effect direction of the significant variants from the MR-MEGA meta-analysis.

Description: Number of significant SNPs per chromosome with effect direction identified by MR-MEGA. The observed p value significance threshold is p < 5×10^-9^. Traits included were (A) stature (height at sacral bone), (B) fore udder position, (C) udder central ligament and (D) front teat position.

**Additional file 18 Figure S17**

Format: png

Title: Association testing for front teat position in HOL cattle with known POLLED genotype.

Description: The upper panel shows the results of an association study between imputed sequence variant genotypes on BTA1 and front teat position in 3,397 HOL cows with known POLLED genotype. The lower panel shows the results of an association study between imputed sequence variant genotypes on BTA1 and front teat position in 3,073 HOL cows not carrying the Friesian polled variant. The red symbol represents the most significantly associated marker from the across-breed meta-analysis

