## Supplementary figures and images for "Across-breed analyses of genome-wide association studies for stature and mammary gland morphology in cattle reveal pleiotropic effects of the Friesian POLLED haplotype"

### Additional File 2 Supplementary Figure 1.png

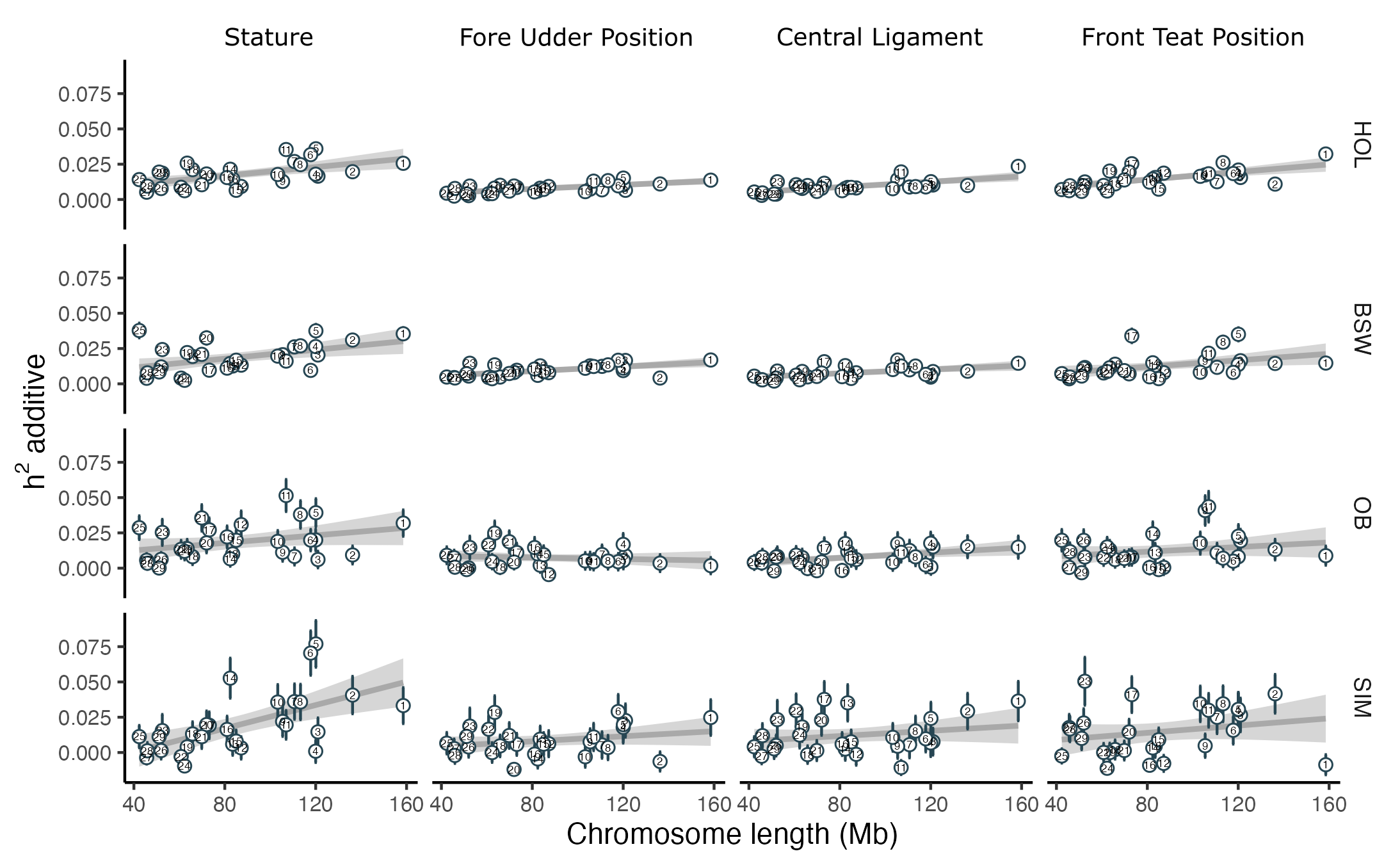

### Additional File 3 Supplementary Figure 2.png

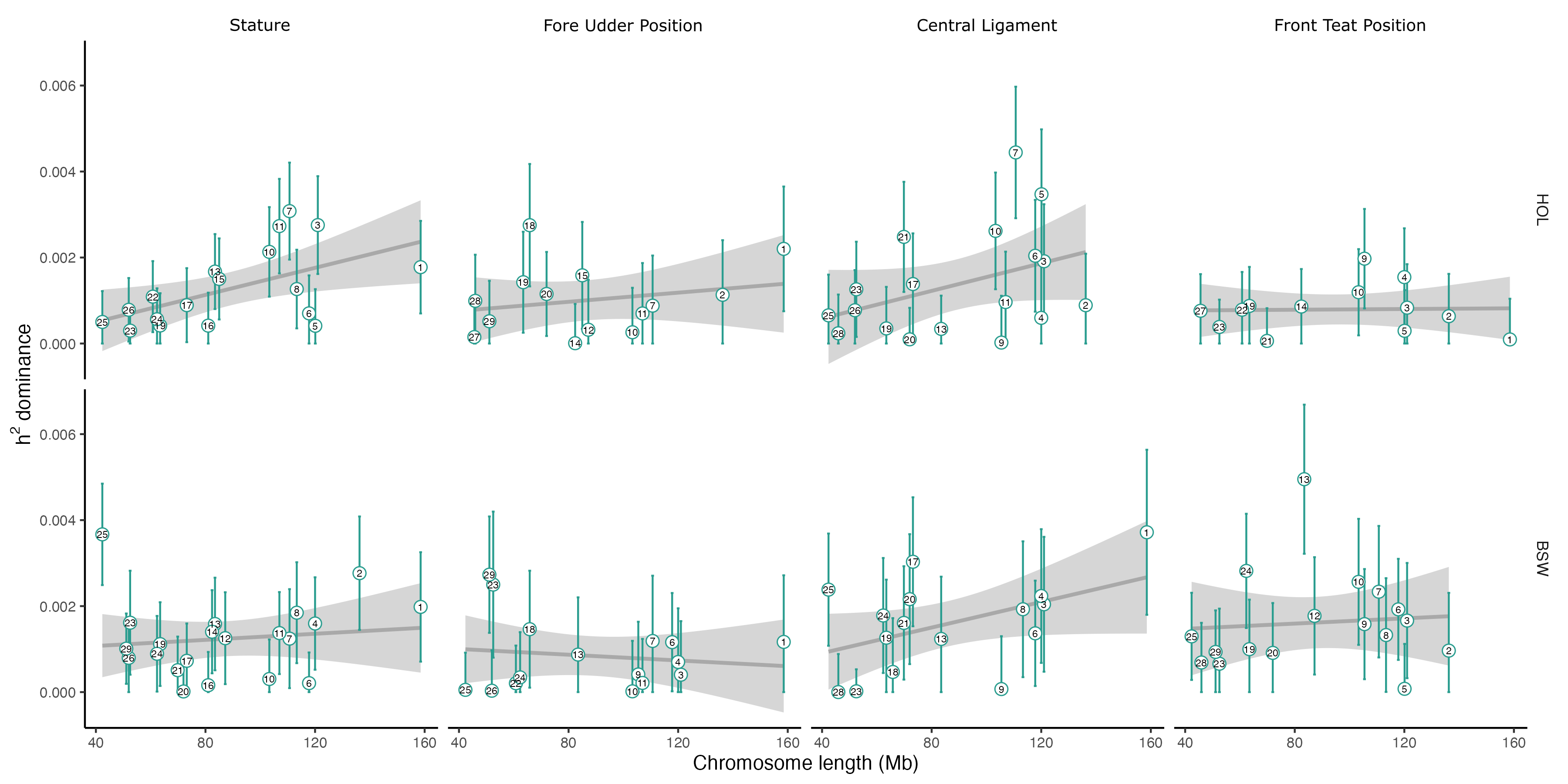

### Additional File 4 Supplementary Figure 3.pdf

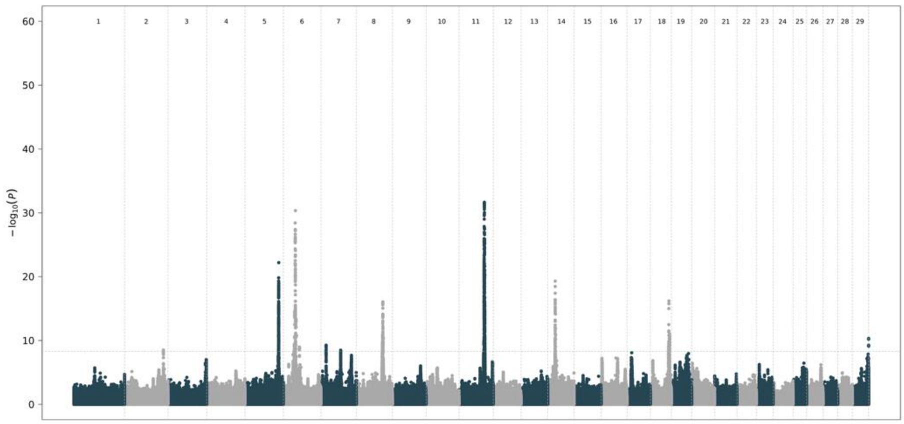


HOL

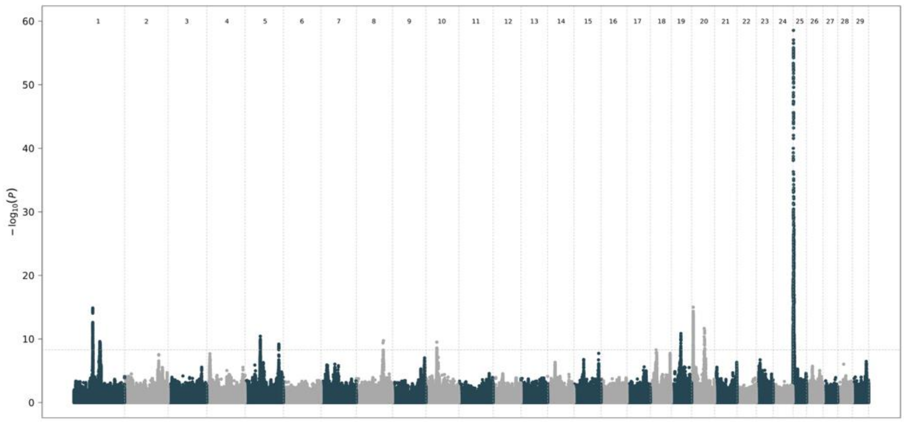


BSW

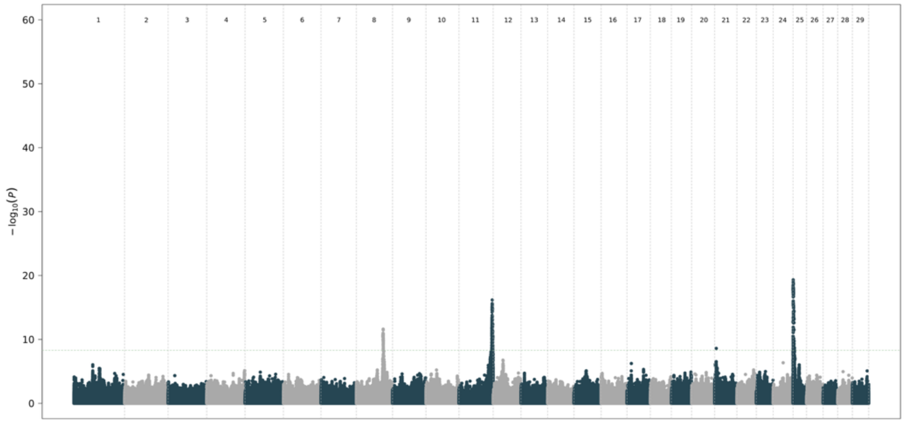


OB

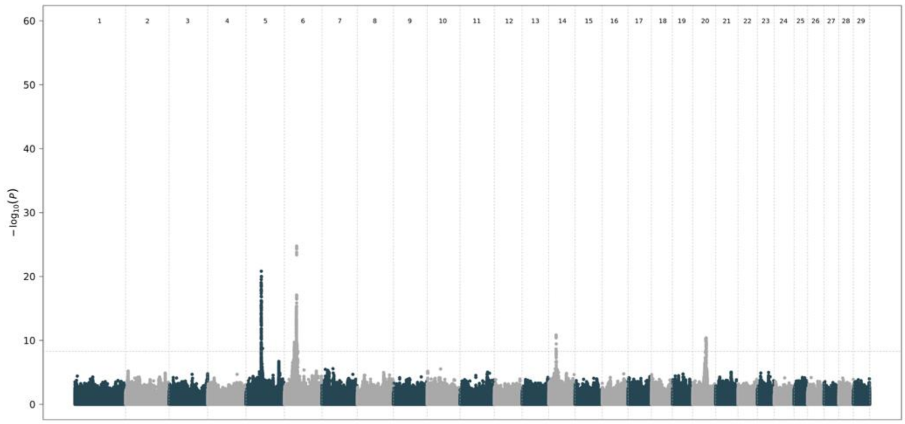


SIM

### Additional File 5 Supplementary Figure 4.pdf

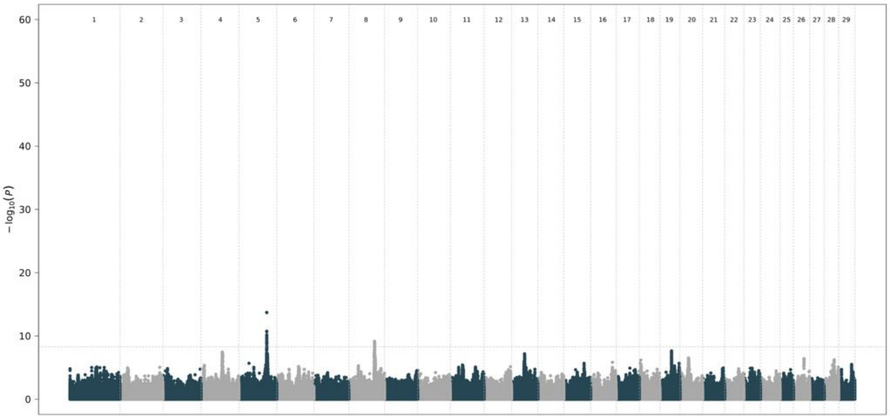


HOL

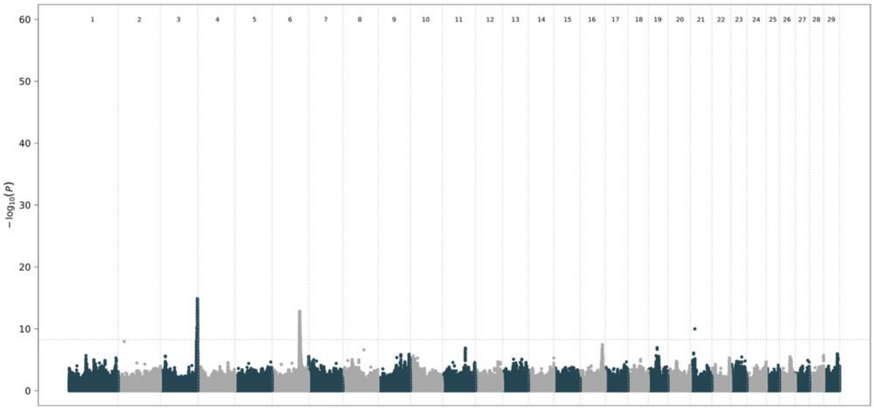


BSW

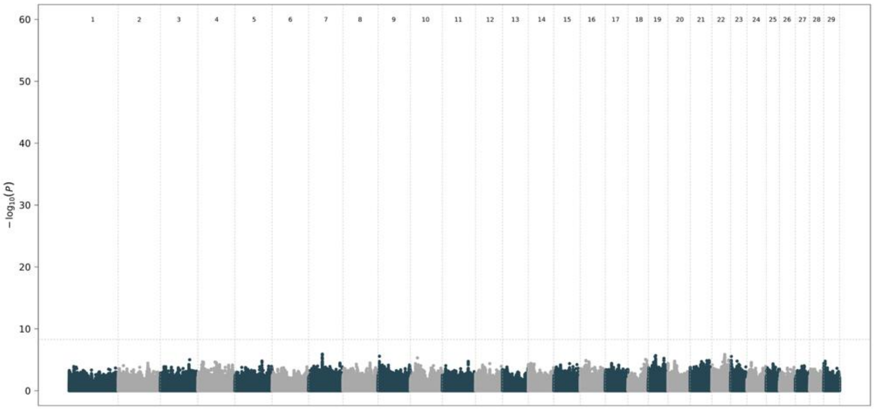


OB

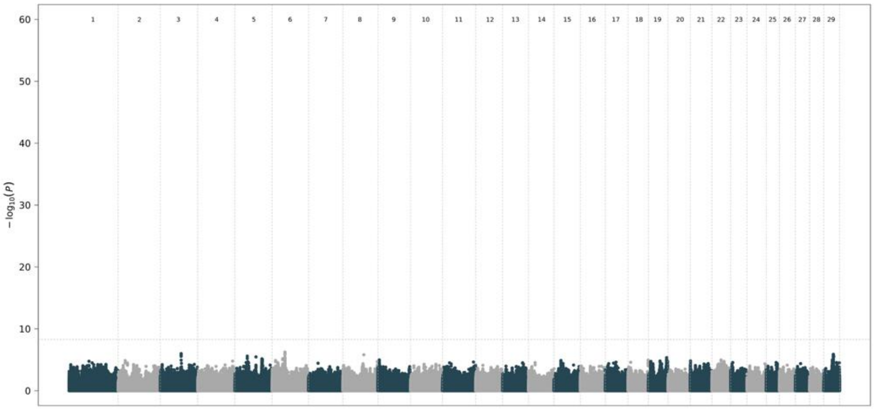


SIM

### Additional File 6 Supplementary Figure 5.pdf

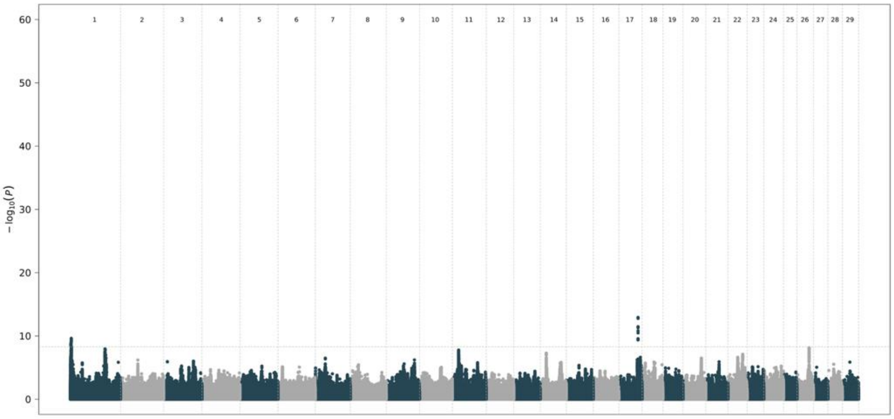


HOL

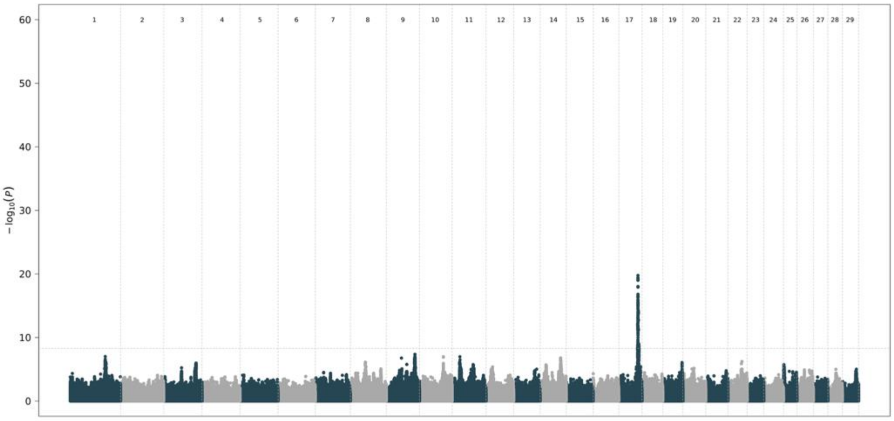


BSW

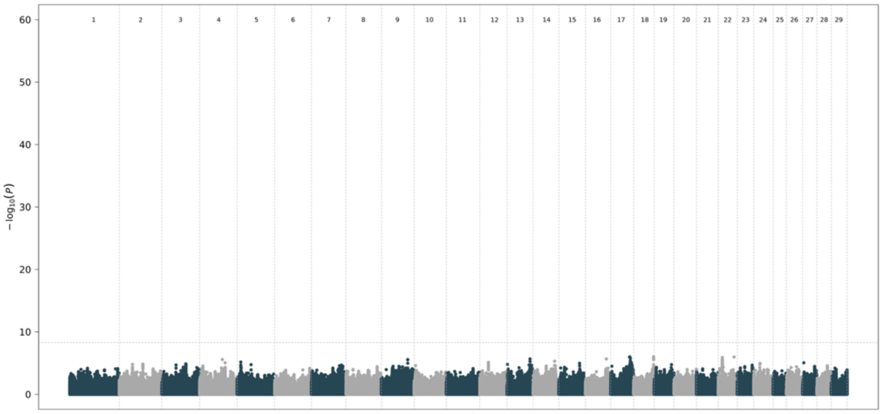


OB

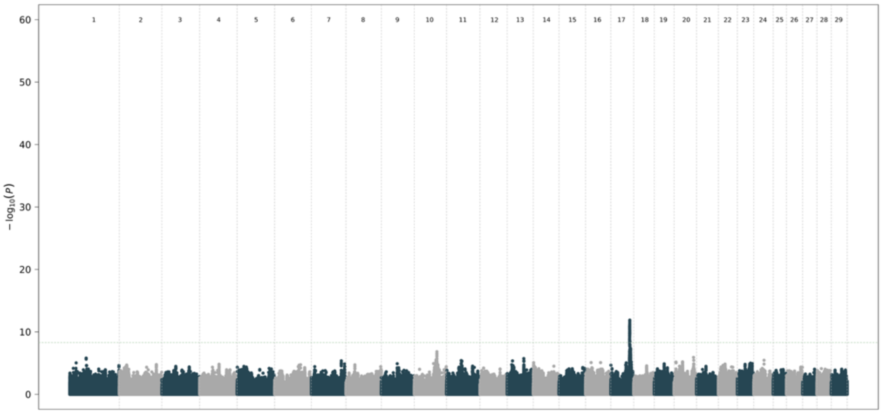


SIM

### Additional File 7 Supplementary Figure 6.pdf

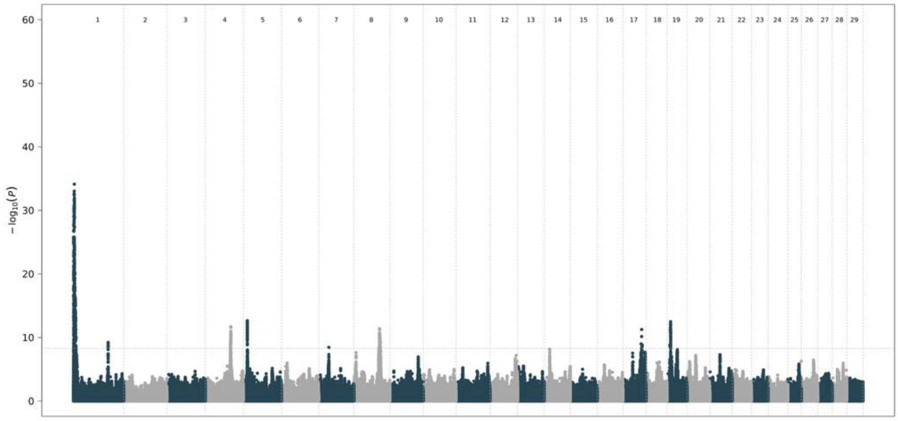


HOL

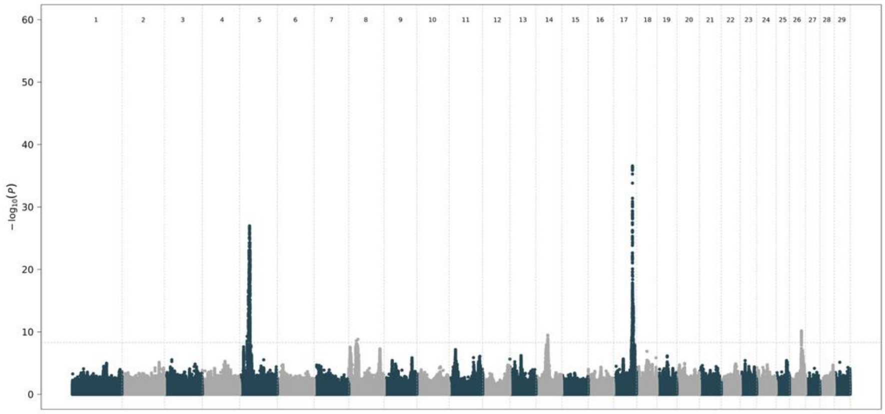


BSW

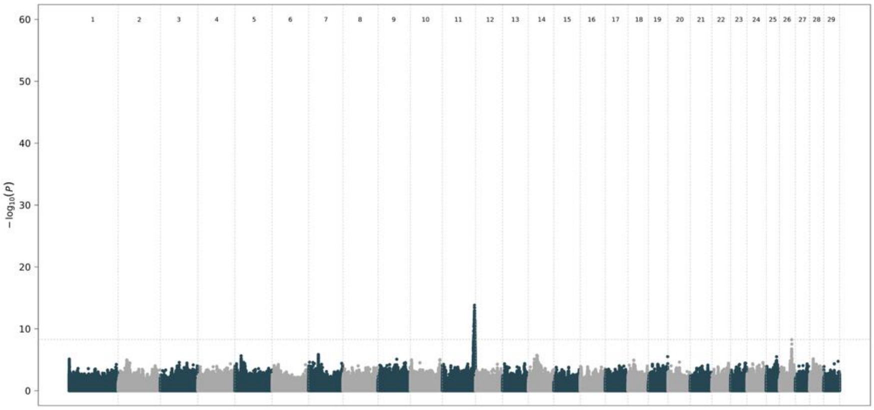


OB

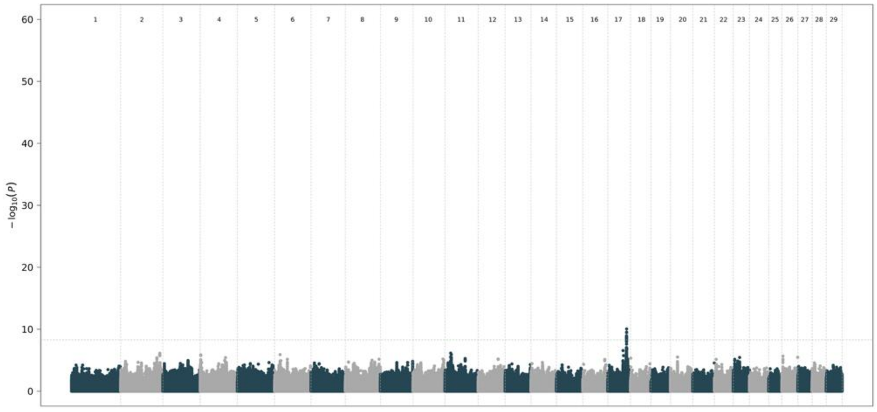


SIM

### Additional File 8 Supplementary Figure 7.pdf

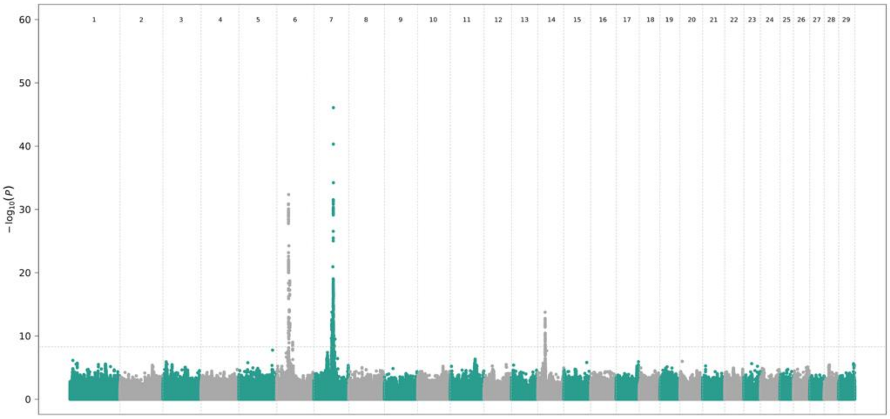


HOL

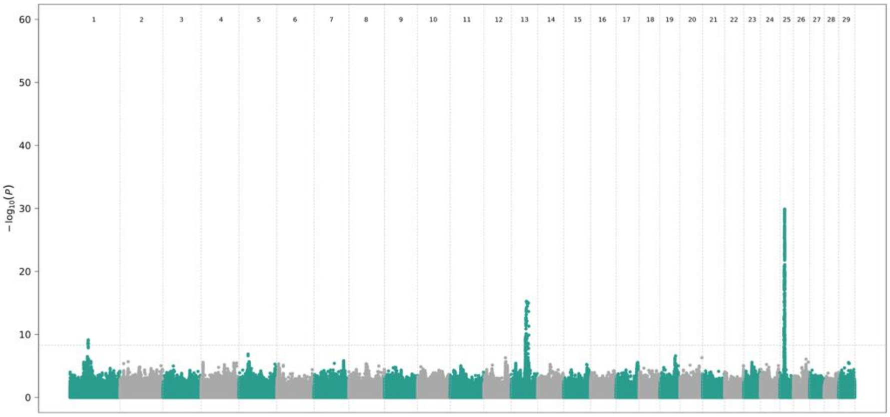


BSW

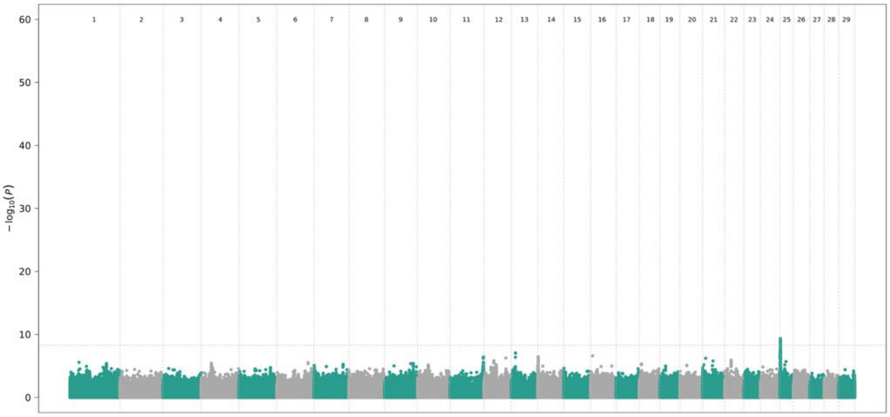


OB

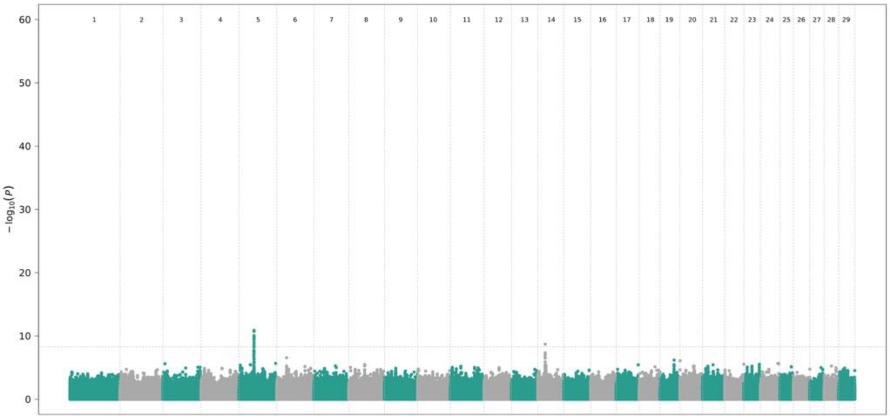


SIM

### Additional File 9 Supplementary Figure 8.pdf

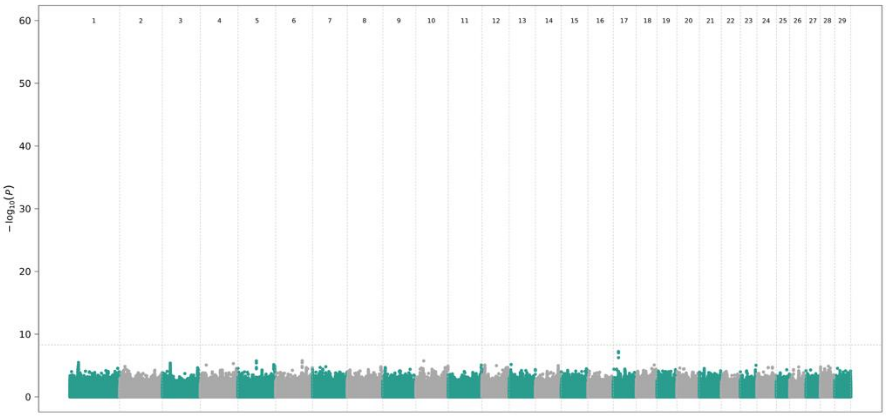


HOL

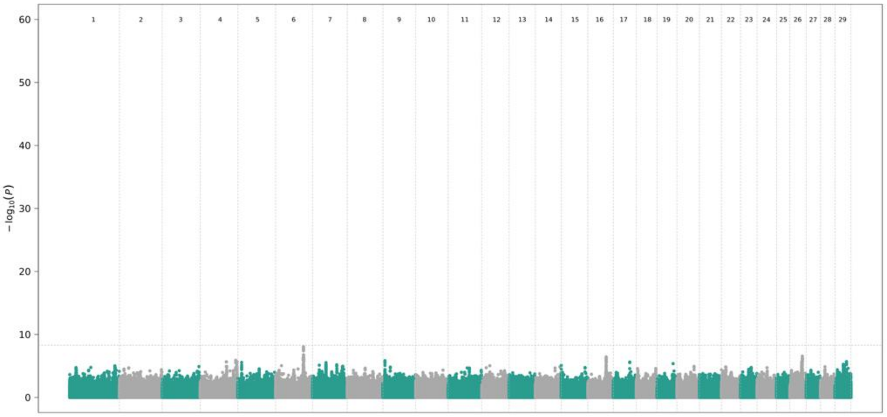


BSW

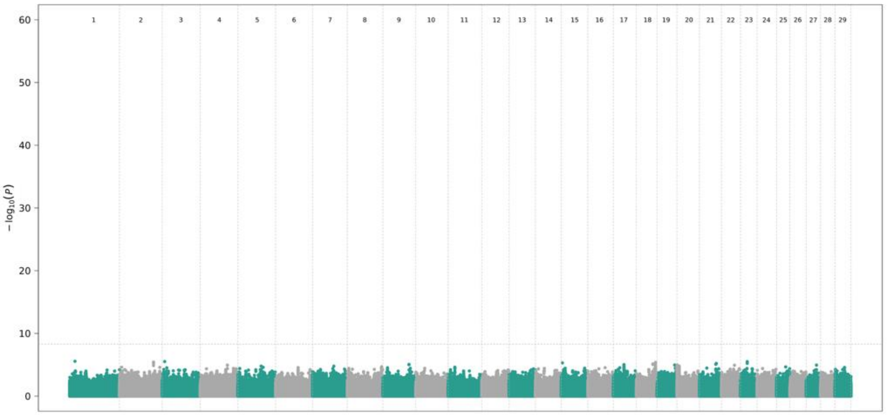


OB

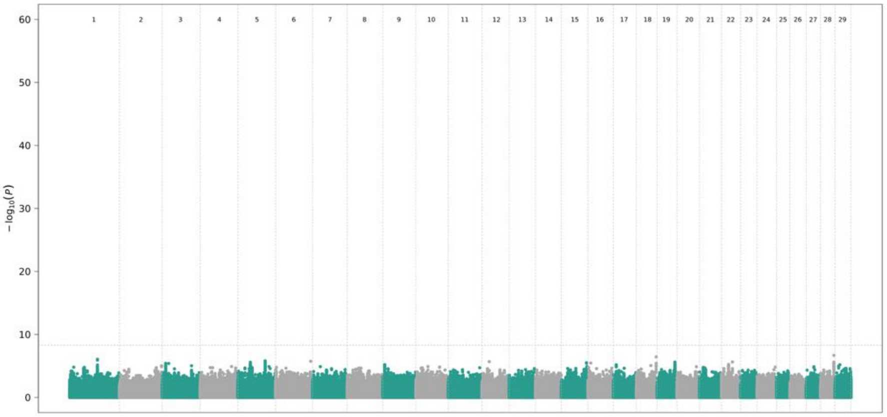


SIM

### Additional File 10 Supplementary Figure 9.pdf

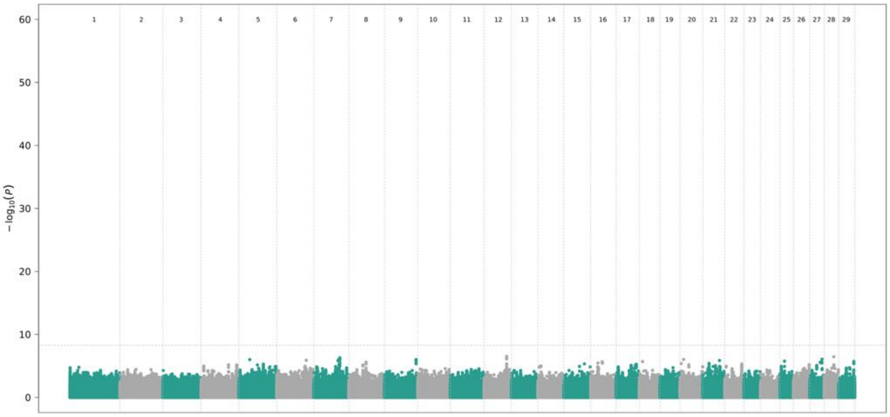


HOL

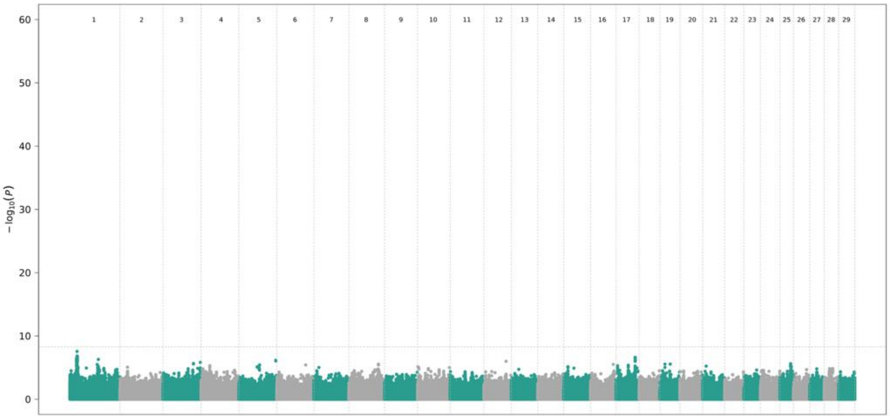


BSW

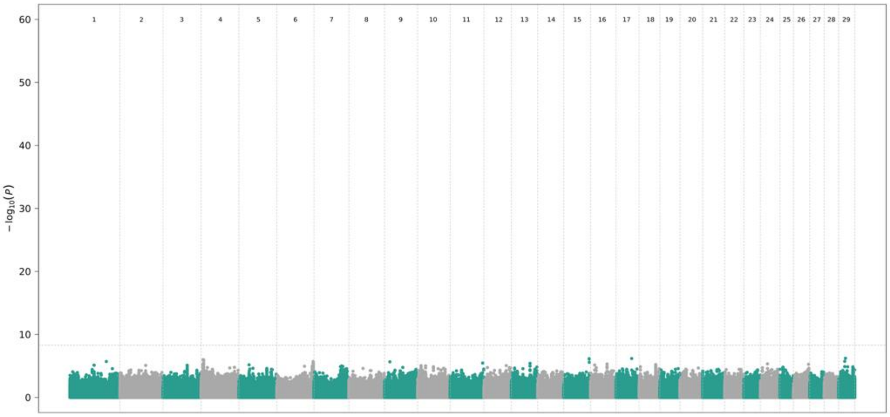


OB

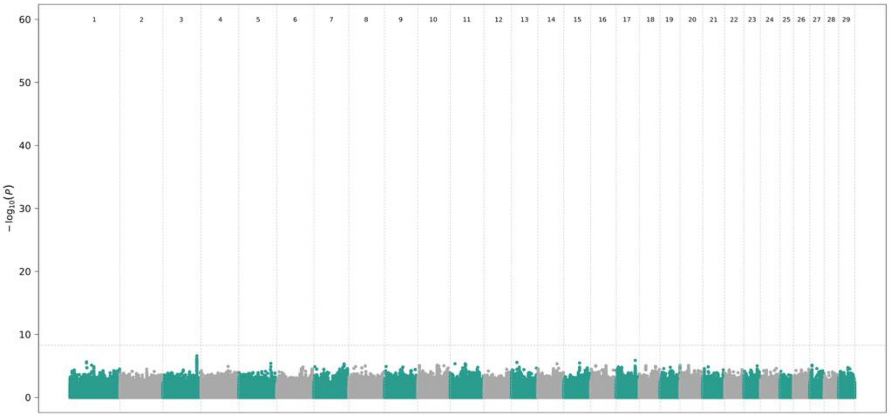


SIM

### Additional File 11 Supplementary Figure 10.pdf

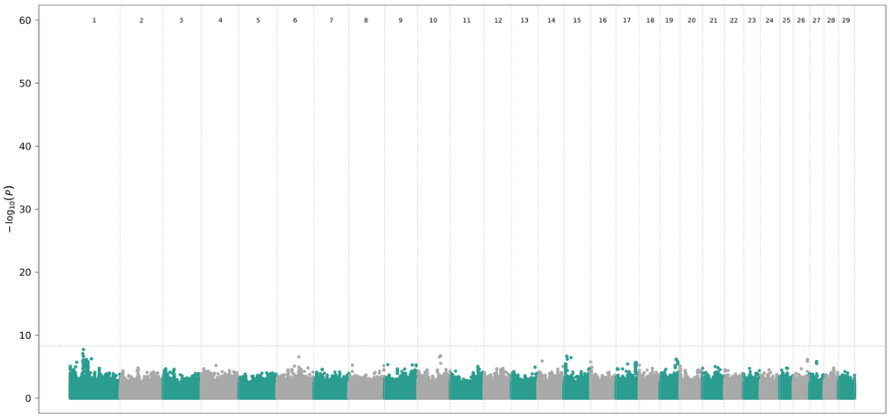


HOL

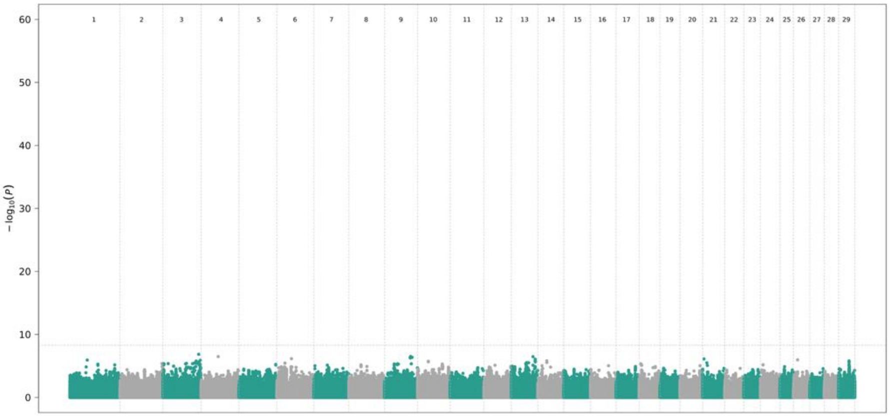


BSW

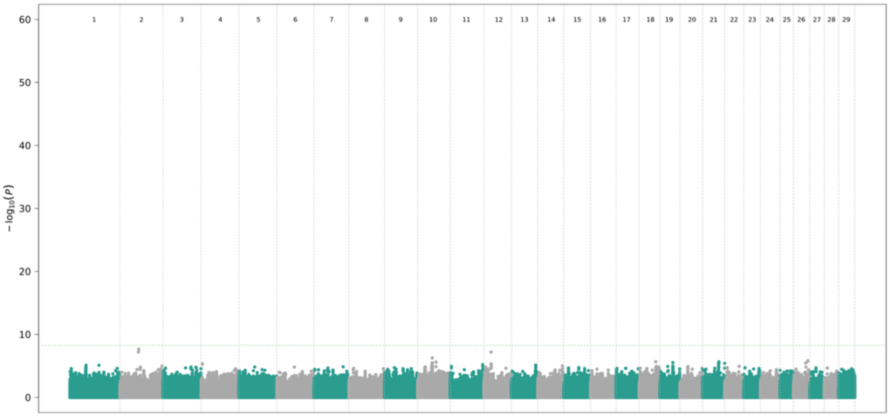


OB

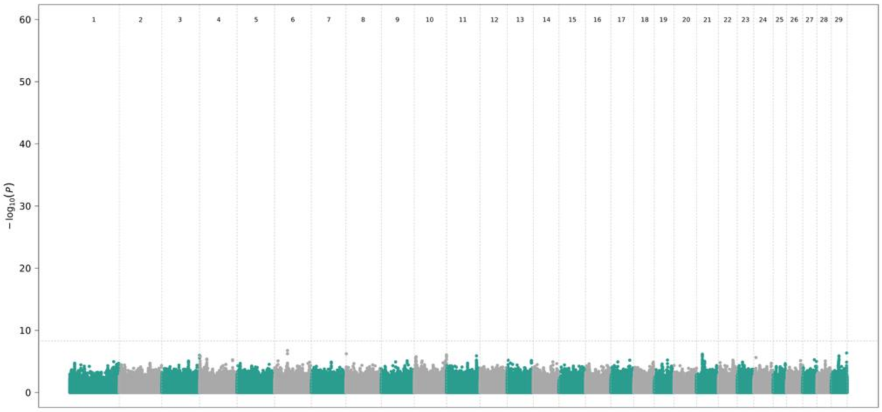


SIM

### Additional File 12 Figure S11.pdf

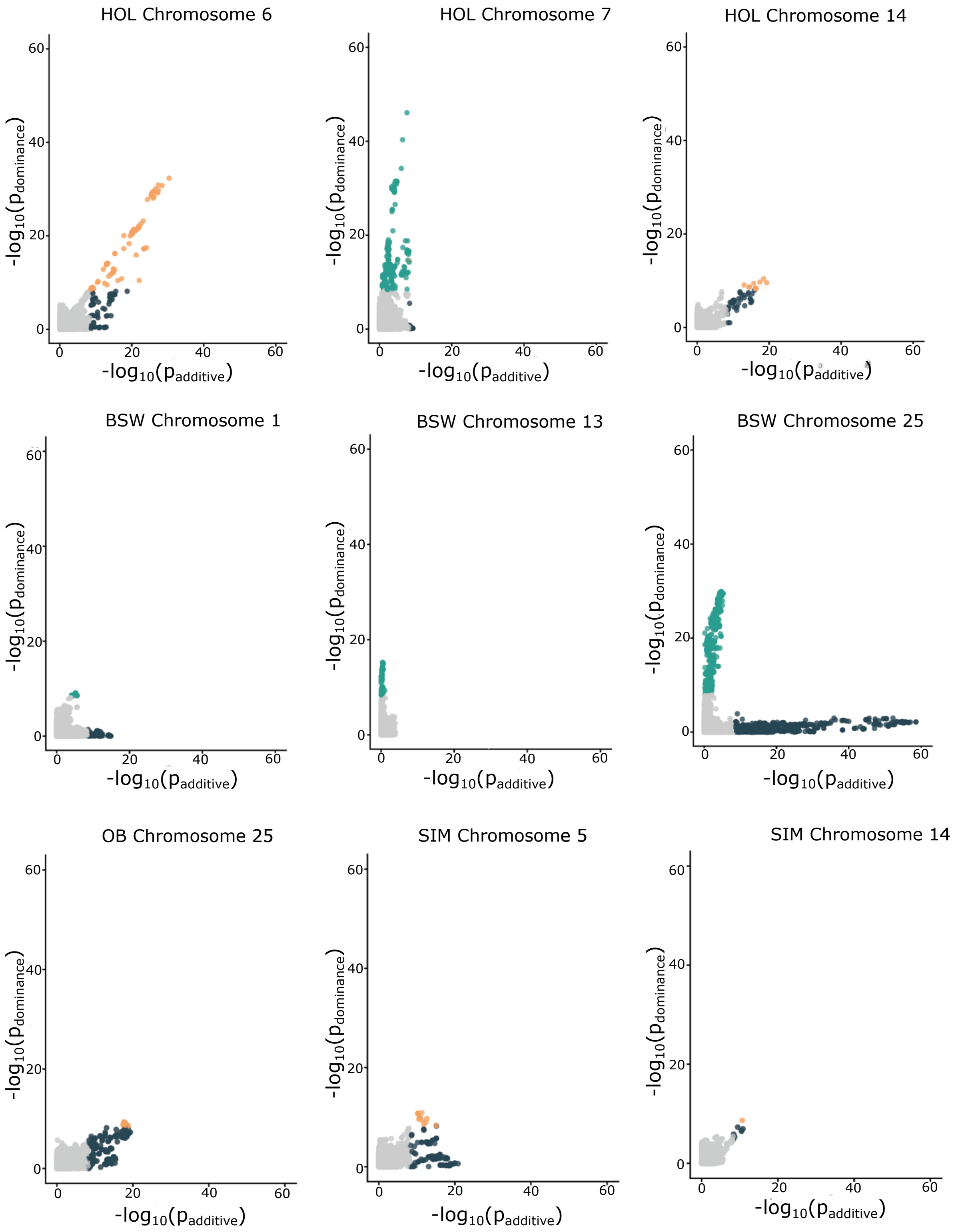

### Additional File 13 Supplementary Figure 12.pdf

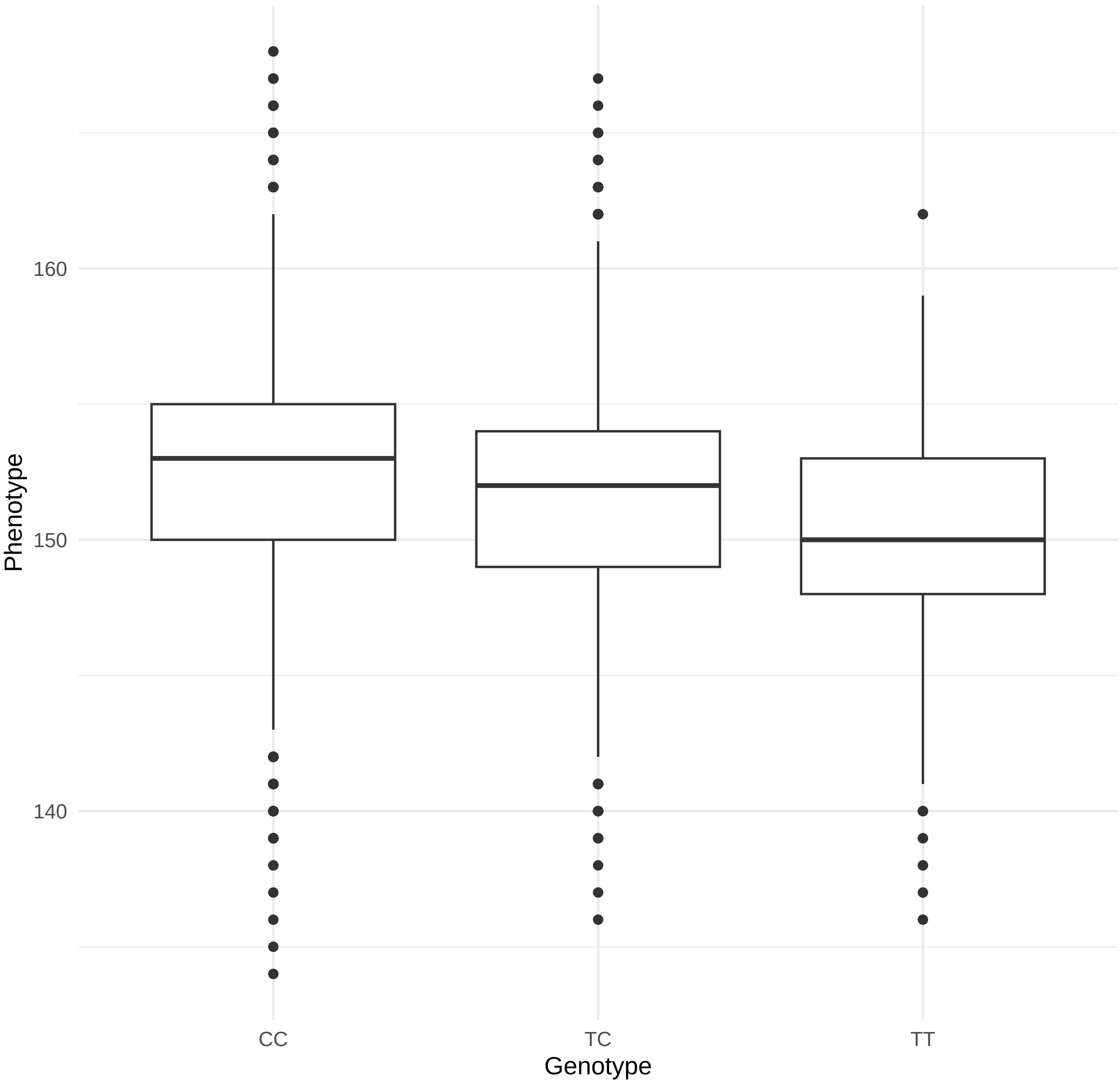

### Additional File 14 Supplementary Figure 13.pdf

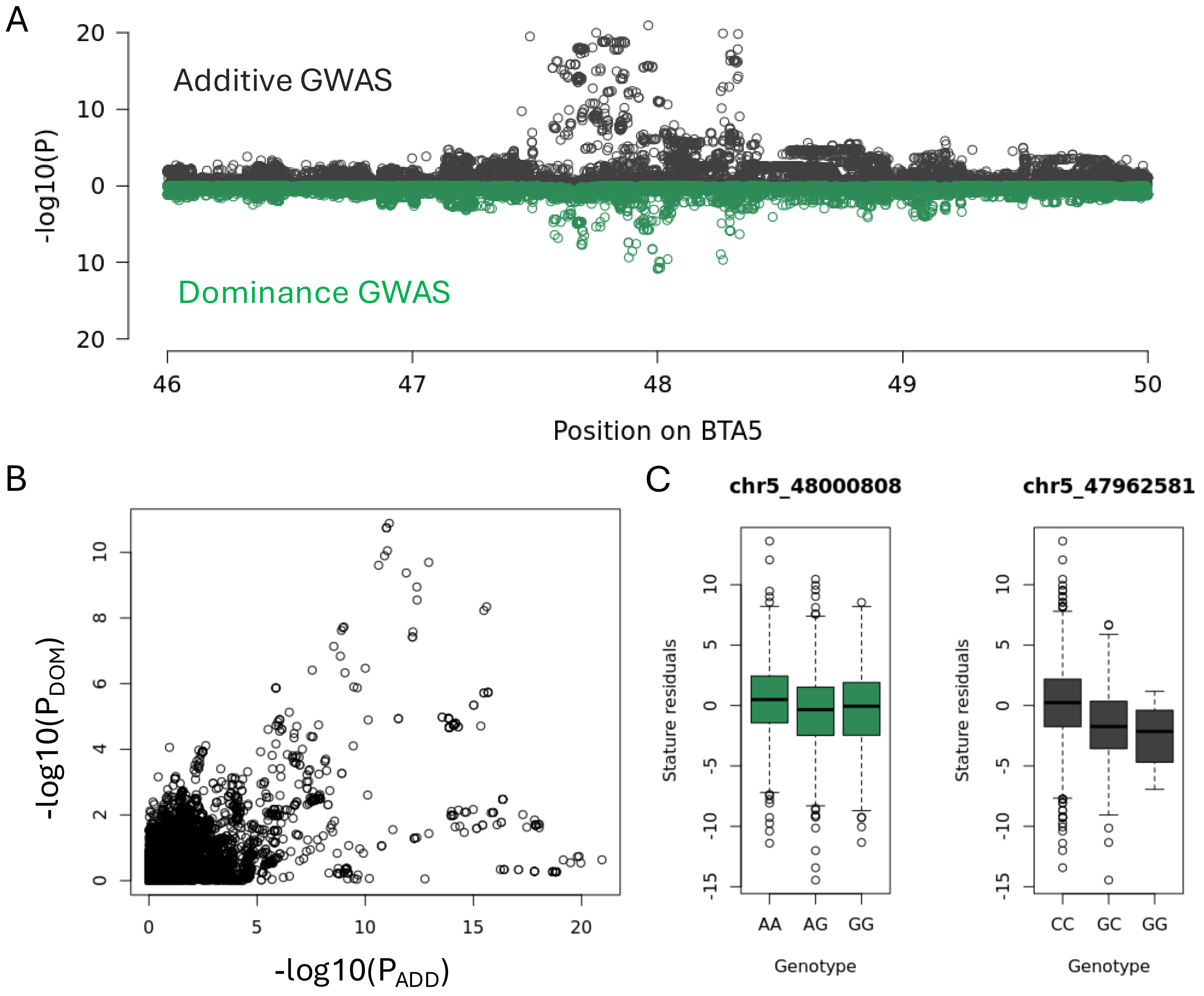

### Additional File 15 Supplementary Figure 14.pdf

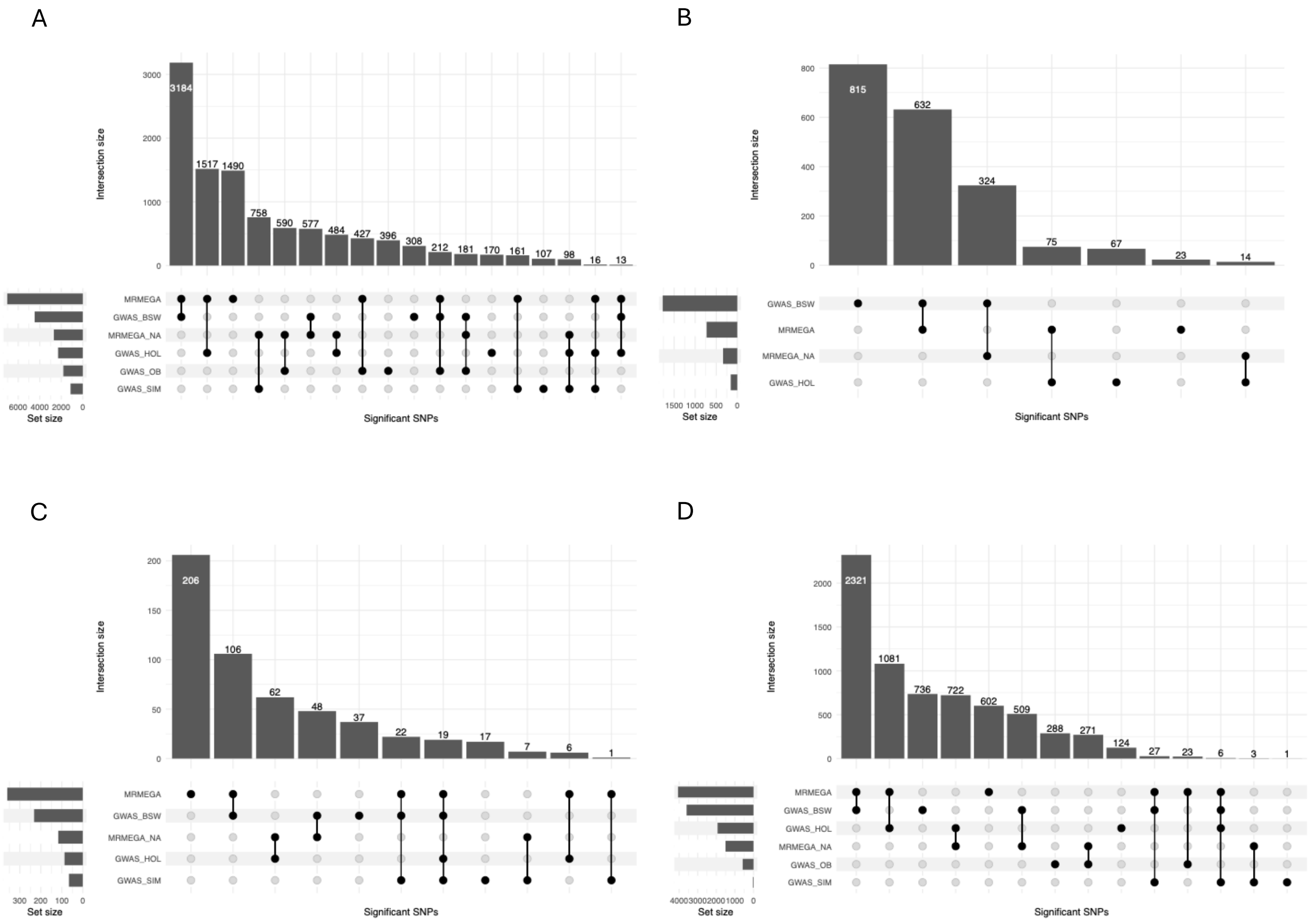

### Additional File 16 Supplementary Figure 15.pdf

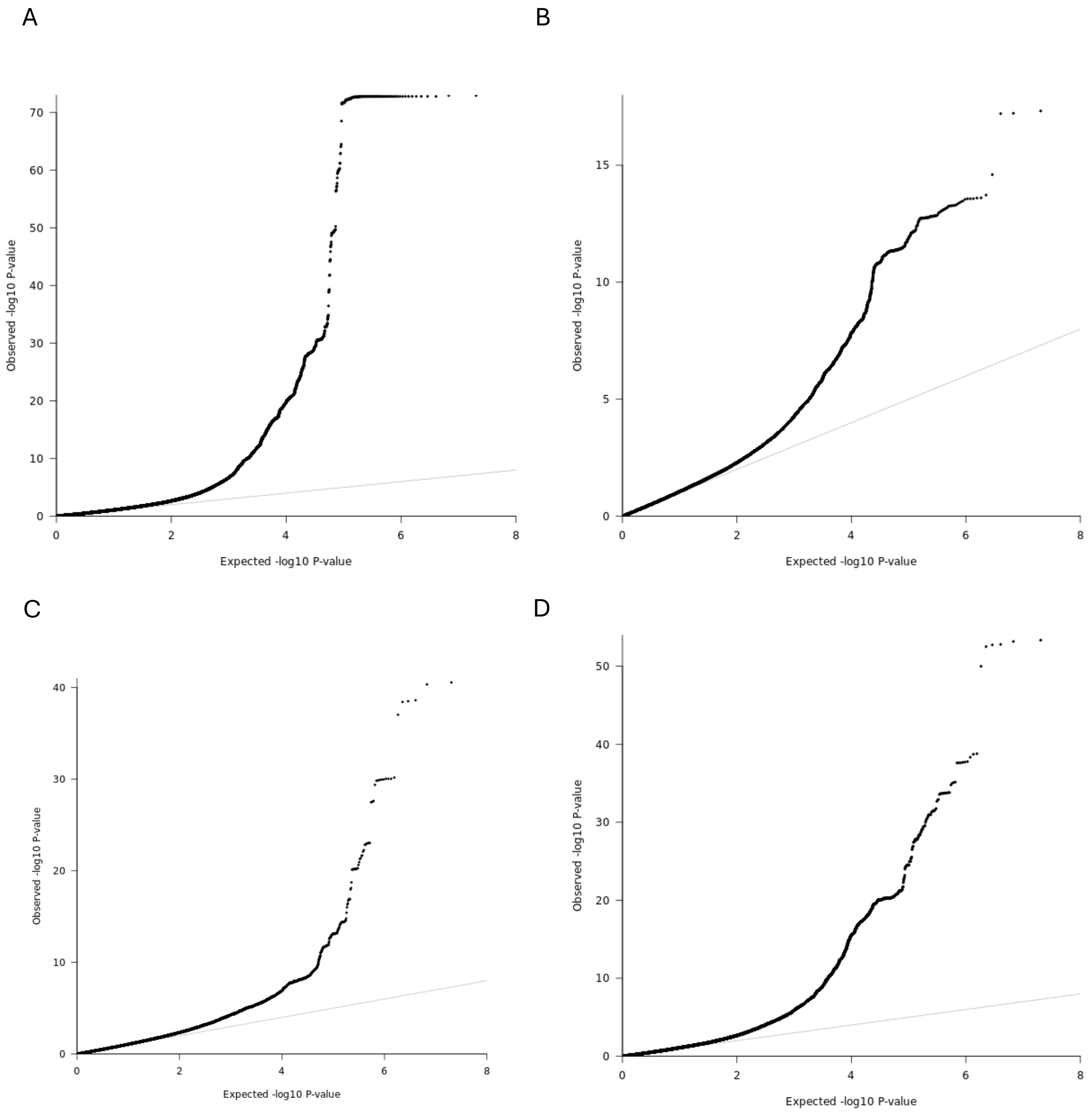

### Additional File 17 Supplementary Figure 16.pdf

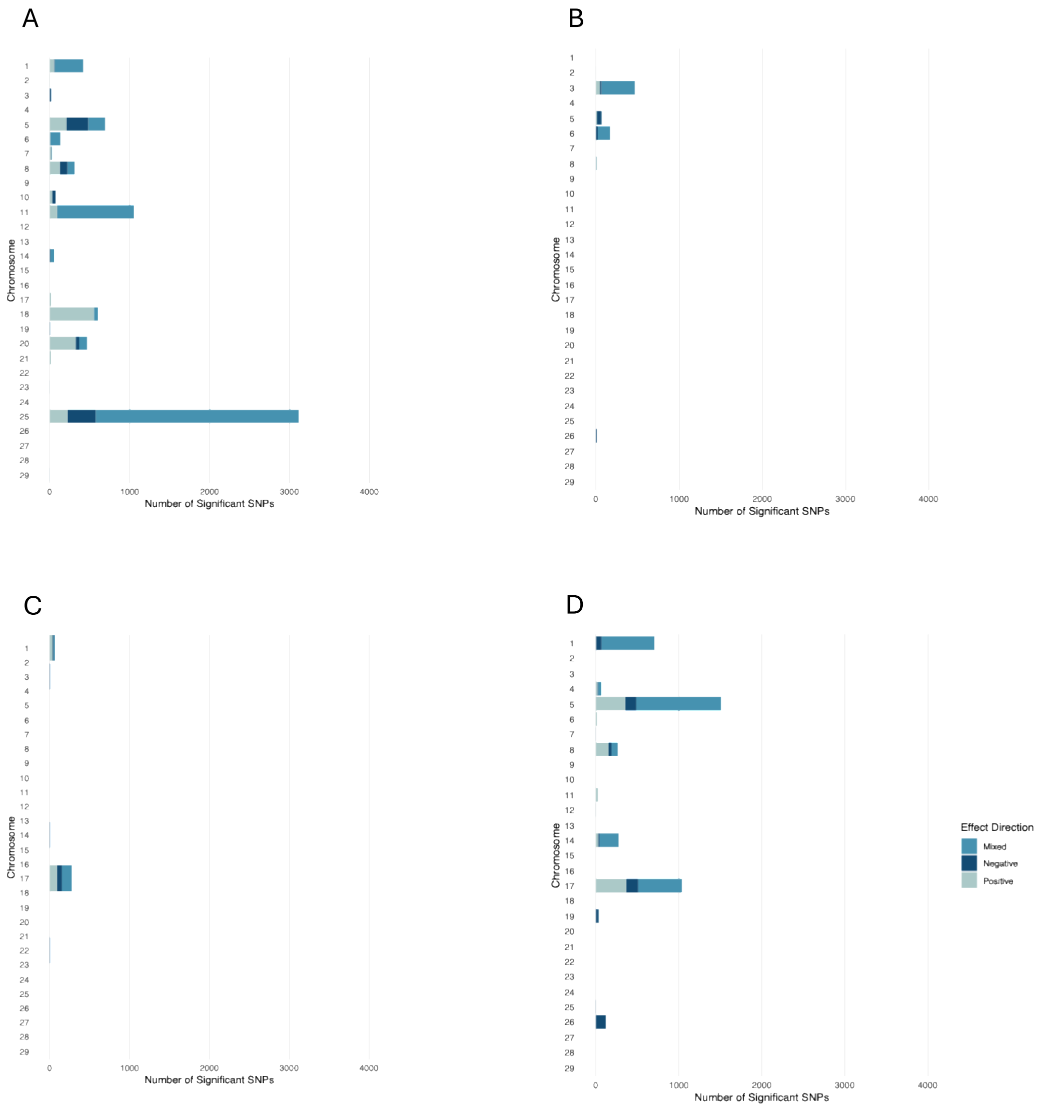

### Additional File 18 Supplementary Figure 17.png

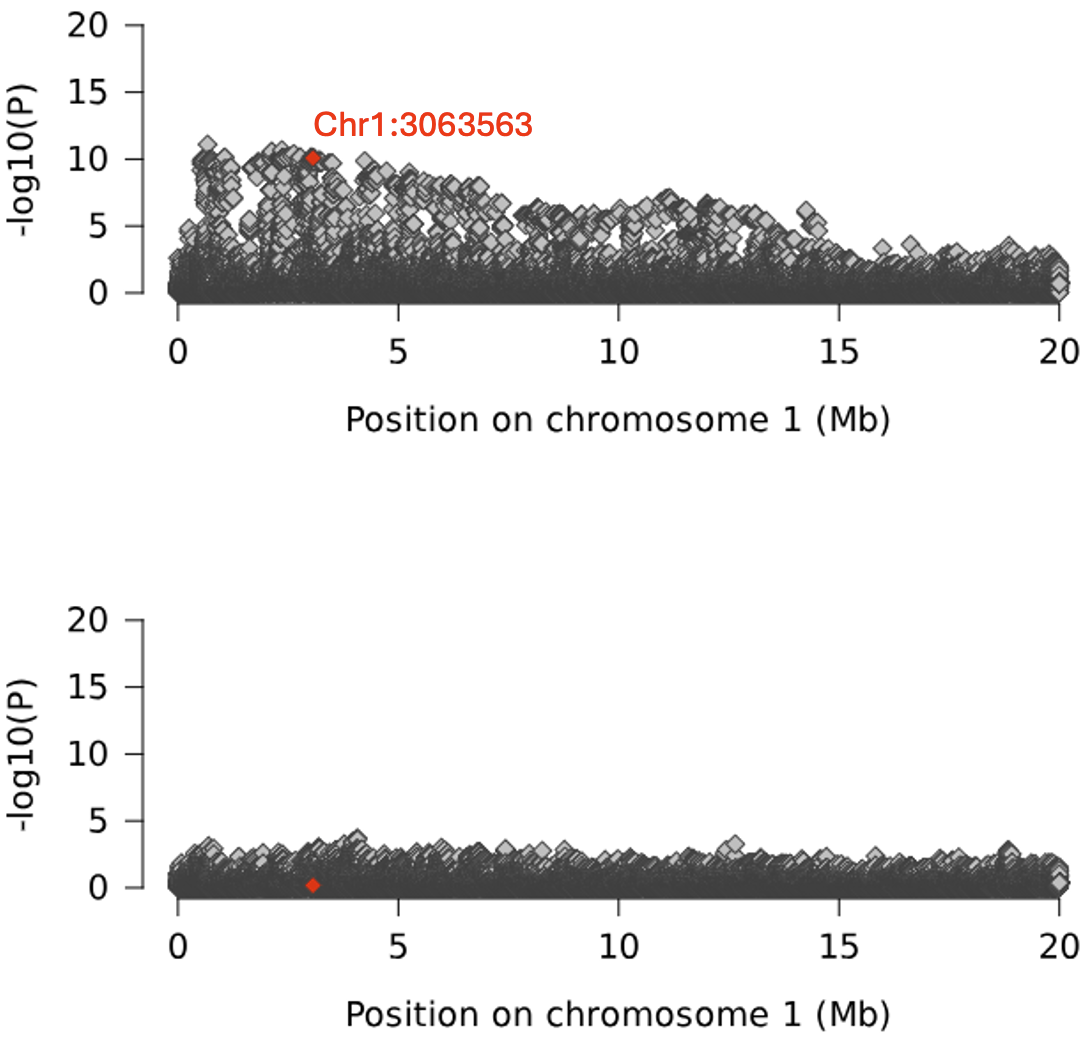
